# A Quantitative Approach for Assessing Multidrug Resistance in Cancer

**DOI:** 10.1101/2024.11.02.621654

**Authors:** Ashley E. Ray, Eugene F. Douglass

## Abstract

Multidrug resistance (MDR) is a primary barrier to successful cancer treatment with small molecule drugs. One mechanism of resistance is through transporters which pump drug out of cancer cells before they exert any therapeutic effect. The most studied transporter is P-glycoprotein (Pgp), a member of the ATP Binding Cassette (ABC) superfamily of drug pumps. Prior research has focused on determining whether drugs are substrates of Pgp. Pgp specificity for FDA approved drugs is currently unclear due to technical variability in assays that quantify enzyme kinetics using non-cellular experimental models. Unfortunately, there is extreme variability in enzyme parameters for drugs that are characterized and gaps in literature for drugs that have not been characterized. Thus, our overall goal is to develop live cell methods to obtain quantitative scores for substrates in cells and fill gaps in the literature. Studying Pgp in the context of MDR, we improve Pgp specificity scores by leveraging new Pgp expression (cell lines, tissues) and function (drug screening) datasets. We experimentally and computationally integrate functional dataset information to better understand Pgp specificity using an approach based on underlying Michaelis-Menten enzyme kinetics. We obtain consensus scores for Pgp specificity across ∼150 FDA approved oncology drugs and validate them experimentally in a subset of 76 substrates selected to represent the spectrum of drugs for Pgp specificity. These scores can be used to calibrate clinical diagnostics (Pgp expression), and our experimental platform can be used to quantify Pgp function in clinical samples. Overall, we develop a parallel computational and experimental procedure to estimate Pgp selectivity in live cells. This procedure can be expanded to other drug transporters which contribute to MDR to further characterize this phenotype quantitatively.

**Significance Statement:** MDR is facilitated through the action of drug transporters which are upregulated in cancer. Even though selective inhibitors have been designed to decrease drug efflux, they failed clinically. So, current clinical practice is to select non-substrates. However, non-substrates are difficult to define. Here, we have developed a quantitative platform for characterizing MDR in cancer that factors in gene expression and enzyme kinetics. Our computational and experimental platform successfully identified the spectrum of substrates for Pgp from the FDA approved oncology drug library. This quantitative approach reflects the multigene phenotype of cancer drug resistance and can be applied to other MDR transporter genes to optimize drug selection clinically.

## Introduction

Drug response in cells is complex. Back in the 1800s, Paul Ehrlich worked on the chemical composition of drugs and their mechanism of action (Ehrlich et. al., 1957). He worked on anti-toxins and wanted to determine how to make anti-toxins 100% neutralizing to the toxins. Ehrlich studied the idea of complementarity where an anti-toxin binds to the target toxin but not to anything else. This led Ehrlich to study enzyme-substrate affinity and gave way to his work on receptors. He then proposed the ‘lock and key’ model for receptors which linked the chemical structure of compounds to pharmacological activity. Essentially, the lock is the enzyme, and its substrate is the key which fits into a specific binding pocket within the enzyme. This fit enables the compound’s pharmacological activity (Bosch et. al., 2008).

Ehrlich proposed the concept of receptor theory, but Archibald Hill made it quantitative. In the early 1900s, Archibald Hill introduced the Hill equation to describe the interaction between hemoglobin and oxygen (Hill, 1910). After one oxygen binds to hemoglobin, it becomes easier for more oxygen molecules to bind other sites on the hemoglobin as the active sites change conformation to increase the likelihood of additional binding. This is depicted by the Hill equation and describes how drugs bind to their targets: % binding = [Drug] / ([Drug] + Kd). As the intracellular binding site becomes saturated with substrate, the curve for binding produces a sigmoidal shape (Gesztelyi et. al., 2012).

The Hill equation models a drug binding directly to its target protein with the dissociation rate constant, Kd, as a measure of binding affinity. Once an enzyme is saturated, it leads to dose response curves where the excess substrate is bound to all available enzymes. In cells, the % binding can be approximated to the effective concentration 50 (EC50) which measures the concentration of drug needed to achieve 50% cell death (Fig S1). In the Hill equation, this assumes that all drug inside the cell becomes bound to the intracellular target.

While the Hill equation works well for protein-based binding assays, it is incomplete for cells. When the Hill equation is applied to cells, there are several implicit assumptions made which can be inaccurate. One key assumption is that the drug concentration inside the cell equals the drug concentration outside the cell ([drug]in = [drug]out), so the EC50 can be approximated to the Kd meaning all substrates bound to the target are linked to its effect. In most cases, this is true, but enzymes such as drug pumps change the equilibrium, so the [drug]in << [drug]out.

Xenobiotic enzymes defend the body from diverse toxins and therefore, do not fit the ‘lock and key’ concept of substrate specificity. They help the human body eliminate, detoxify and metabolize drugs. Thus, most xenobiotic enzymes are expressed in the liver, intestines, kidneys, lungs, blood and brain. The largest class of xenobiotic enzymes is the Cytochrome P450 (CYP450) family which are mainly involved in the first phase of metabolism (Omura et. al., 1962). The goal of Phase I drug metabolism is to create a polar, water soluble metabolite through oxidation, reduction or hydrolysis. The product of Phase I leads to Phase II drug metabolism in which metabolites are conjugated to charged species through glucuronidation, sulfation, glutathione conjugation, methylation and acetylation. The addition of these ionized groups makes the metabolites more water soluble and higher molecular weight, so they are more easily excreted from the body (Wrighton et. al, 1992, McDonnell et. al., 2013).

However, CYP450 enzymes have been implicated for their role in multidrug resistance (MDR). CYP450 enzymes are not only expressed in tissues but within individual cells. In cells, they metabolize a fraction of drug when it enters the cell. This is a protection mechanism to prevent high levels of drug from accumulating and causing toxicity (Meijerman et. al., 2008, Runge et. al., 2000, Benet et. al., 1999).

Additionally, a third group of xenobiotic enzymes are the drug transporters. These enzymes also protect the tissues and cells by pumping out drug. Drug transporters have an important physiological role and are highly expressed in barrier tissues and tissues involved in drug pharmacokinetics in the body. Tissues with high amounts of drug transporters include the brain, intestines, colon, liver, kidneys and testes (Fig S2) (Benet et. al., 1999, Faber et. al., 2003, Oostendorp et. al., 2009, Inui et. al., 2000, Su et. al., 2009, Ebinger et. al., 2006).

The most notable group of drug transporters are the ATPase Binding Cassette (ABC) transporters (Gottesman et. al., 2001). ABC transporters encompass subfamilies of which ABC subfamilies B, C and G are best known for conferring drug resistance. ABCB1 is the gene for Permeable glycoprotein (Pgp), an extensively studied drug transporter in the context of cancer MDR (Szakacs et. al., 2006).

Pgp defies standard pharmacologic theory because it pumps out various substrates (Ploeger et. al., 2009). This promiscuity can be traced to Pgp’s globular binding pocket which can accommodate many different chemical structures of drugs. MDR enzymes typically follow pseudo-1^st^ order reaction kinetics (Southwood et. al., 2018). Since Pgp can pump out many different substrates, it has a large Km because its affinity is not very high for any one substrate. Thus, we can assume that the [drug]in is much lower than the Km which means that we can approximate the EC50 to the Km in the linear regime of the reaction.

As a promiscuous enzyme, Pgp has been described as a vacuum cleaner by “sucking up” drug from the cell membrane and pumping it out (Raviv et. al., 1990). Pgp is the first ABC transporter discovered and accordingly, the drug pump most studied in the scientific literature (Knox et. al., 2011). As a family, ABC transporters are promiscuous and accommodate many drugs though they have preference for different substrates which underlies their physiological role. The description of drug pumps as “vacuum cleaners” is perhaps a more accurate depiction as they pump out a variety of drugs and biochemical molecules (Raviv et. al., 1990). This contrasts with Ehrlich’s description of enzyme receptors through a ‘lock and key’ model where one substrate fits into one enzyme (Ehrlich et. al., 1957).

In the 1900s, Michaelis-Menten enzyme kinetics came from the study of a reaction yielding fructose and glucose from sucrose catalyzed by invertase (Michaelis et. al., 1913). For this reaction, Michaelis and Menten discovered the formation of an enzyme-substrate complex by measuring the velocity of the reaction as a function of sucrose concentration. Michaelis-Menten enzyme kinetics are different from the Hill equation in that they incorporate reaction velocity as a function of enzyme concentration and substrate turnover (Johnson et. al., 2011).

The output of the Hill equation is a measure of binding affinity of a substrate for an enzyme (Equation 1). The Michaelis-Menten equation expands on the Hill equation by changing the Kd to Km and measuring reaction velocity (Equation 2). The Km is the substrate concentration at which the enzyme reaction is half-maximal, so a smaller Km indicates greater binding affinity. The Michaelis-Menten equation defines the maximum velocity, Vmax of the enzyme-catalyzed reaction as a function of enzyme concentration, [E] and rate of substrate turnover, kcat (Equation 3). Since a greater enzyme concentration and substrate turnover increases the maximum reaction velocity, these factors contribute to an overall greater reaction velocity (Equation 4).

Equation 1: % binding = [Drug] / ([Drug] + Kd)

Equation 2: velocity =([E] ^*^ kcat ^*^ [Drug]) / ([Drug] + Km)

Equation 3: Vmax = [E] ^*^ kcat

Equation 4: velocity = (Vmax ^*^ [Drug]) / ([Drug] + Km)

As with the Hill equation, there are some key assumptions made in the context of modeling MDR with drug transporters. First, the contribution of enzymes to MDR is additive with each enzyme contributing independently to flux. Second, Pgp is a promiscuous enzyme with a high Km for multiple drugs, so the Km is much larger than the [drug]in. Third, drug influx is impacted by diffusion only, so [drug]in = [drug]out at steady state (Michaelis et. al., 1913).

Under 1^st^ order conditions ([substrate] < Km), the membrane transport rate (kefflux) is expected to be the kcat ^*^ [Transporter] / Km (Nikaido, 1994, Mitchell, 1957). This assumption is commonly used for membrane transporters and is consistent with clinical pharmacokinetics where xenobiotic enzymes are assumed to act under 1^st^ order conditions clinically (Southwood et. al., 2018). Thus, Michaelis-Menten enzyme kinetics can be thought of as a linear relationship where y = slope ^*^ x. In this linear relationship, y is the efflux rate (kefflux), slope is (kcat / Km) / kdiff and x is MDR expression ([E]). This corresponds to the change in drug EC50. If we assume pseudo-1^st^ order conditions, then EC50 and [E] are expected to be linear (Fig S3) (Seelig 2007).

This decreased drug potency and resulting EC50 shift is an indication of the MDR phenotype. Fortunately, we have data on this phenotype from new databases such as DrugBank, the Cancer Therapeutics Response Portal (CTRP), Genomics of Drug Sensitivity in Cancer (GDSC), Profiling Relative Inhibition Simultaneously in Mixtures (PRISM) and Cancer Cell Line Encyclopedia (CCLE) (Basu et. al., 2013, Yang et. al., 2013, Corsello et. al., 2020, Ghandi et. al. 2019). To bridge these *in-vitro* databases with clinical data, we examined 34 most FDA-indicated chemotherapies and *in-vitro* data from the Cancer Dependency Map Portal (https://depmap.org/portal, Tsherniak et. al., 2017). Our analysis demonstrated a correlation between biomarkers (direct binding, metabolism, transport, DNA repair) and drug sensitivity (results are shown in more detail below). Further analysis showed that biomarker RNA expression is directly correlated with drug EC50. Thus, a drug’s EC50 change can be modeled by a kinetic ratio in MDR cancers.

There are increasing efforts to digitize raw data from primary literature references which give us the opportunity to compare MDR phenotypes and kinetic parameters across primary literature for the first time. Relevant examples for our quantitative analysis include BRENDA and DrugBank (Schomburg et. al., 2002, 2011, Knox et. al., 2011). Specifically, BRENDA is an enzyme repository for information on enzyme biochemistry, structure, kinetics, function etc. (brenda-enzymes.org/). DrugBank is a database with information on approved drugs including indication, mechanism, pharmacokinetics and pharmacodynamics (go.drugbank.com/). For BRENDA’s goal is to bridge genomics with enzymes. DrugBank provides information on enzymes and chemistry for already approved drugs through the FDA and foreign regulatory agencies.

BRENDA is useful for MDR kinetic parameters such as kcat and Km. The BRaunschweig ENzyme DAtabase (BRENDA) was founded in 1987 at the German National Research Centre for Biotechnology as an enzyme information data system. The need for a systematic collection of enzyme information for genomic interpretation and field application underlies BRENDA. BRENDA was developed to provide more information on gene products and enzymes to match the increasing projects on genome sequencing. When BRENDA was originally published, it contained data from >46,000 primary literature references with data from >40,000 different enzymes. Specifically, BRENDA includes information on enzyme nomenclature, enzyme structure, enzyme-ligand interactions, functional parameters, molecular properties, organism-related information and bibliographic data (Schomburg et. al., 2002).

In the 2020s, BRENDA has evolved to include >5 million data from 90,000 enzymes across 13,000 organisms from 157,000 primary literature references (Chang et. al., 2021). Additionally, BRENDA now offers enzyme pathway maps covering metabolic pathways and biochemical processes. Viewers can see chemical reactions and enzyme-ligand information within the pathway maps. Currently, each enzyme has its own Enzyme Summary Page which gives an overview of all available information for it within BRENDA. It also incorporated a new tool which predicts the intracellular localization of each enzyme given its function (Chang et. al., 2021).

## Materials and Methods

### Cell Lines

The ductal adenocarcinoma cell line DU4475 was obtained from the American Type Culture Collection (ATCC). DU4475 cells were maintained in Roswell Park Memorial Institute (RPMI) 1640 medium with GlutaMAX and 25 mM HEPES (Gibco) with 10% fetal bovine serum (FBS) (Fisher Scientific) and 1% penicillin-streptomycin (Gibco) and kept in an CellXpert incubator (Eppendorf) at 37 °C in 5% CO2.

### Inhibitor Screen

DU4475 cells were trypsinized (0.05% Trypsin-EDTA, Gibco), incubated at 37 °C and 5% CO2 for 5 minutes, resuspended in sterile Phosphate Buffered Saline (PBS) (PBS pH 7.4, Gibco) and counted with a hemocytometer and trypan blue (0.4% Trypan Blue Stain, Gibco). 100,000 cells/well and 100 ul of cells/PBS were plated in a black 96 well plate. Ten Pgp inhibitors (SelleckChem) were tested from 0.5 pM to 5 uM. 100 ul of each inhibitor concentration was added to the black 96 well plate. 100 ul of a 1% DMSO (Acros) in PBS solution was added to a subset of wells to serve as negative controls. A staining solution was prepared of 0.36 uM Calcein AM (UltraPure Grade, AnaSpec) in sterile PBS and 100 ul added to all wells. After addition, the fluorescence (Calcein excitation = 485 nm, emission = 526 nm) was measured every 10 minutes for 6 hours at 5% CO2 and 37 °C using a Synergy H1 microplate reader and CO2 controller (BioTek).

### Drug Screen

DU4475 cells were trypsinized (0.05% Trypsin-EDTA, Gibco), incubated at 37 °C and 5% CO2 for 5 minutes, resuspended in sterile PBS (pH 7.4, Gibco) and counted with a hemocytometer and trypan blue (0.4% Trypan Blue Stain, Gibco). 100,000 cells/well and 100 ul of cells/PBS were plated in a black 96 well plate. As before, 100 ul of a 1% DMSO (Acros) in PBS solution was added to a subset of wells to serve as negative controls. 30 ul of each drug (SelleckChem) and 70 ul of PBS were added to each well. A staining solution was prepared of 0.36 uM Calcein AM (UltraPure Grade, AnaSpec) in sterile PBS and 100 ul added to all wells. After addition, the fluorescence (Calcein excitation = 485 nm, emission = 526 nm) was measured every 10 minutes for 6 hours at 5% CO2 and 37 °C using a Synergy H1 microplate reader and CO2 controller (BioTek).

### Flow Cytometry

Quantitative flow cytometry was performed to quantify single cell expression of Pgp on six cell lines (DU4475, DLD1, Caco2, K562, K562-Dox, K562-Imatinib). Cell culture media was removed, and cells taken up in “staining buffer” (Phosphate Buffered Saline (PBS) (pH 7.4, Gibco) + 0.5 mM EDTA (Invitrogen) + 1% BSA (VWR)) to a density of 1 million cells/ml following trypsinization (Caco2 and DLD1) (0.05% Trypsin-EDTA, Gibco) and two washes. 100 ul of each suspension (∼100,000 cells) was added to four different Eppendorf tubes on ice and 4 ul of blocking IgG (Fisher Scientific, P131154) was added and incubated for 5 minutes. 10 ul of phycoerythrin (PE)-conjugated anti-Pgp antibodies (VWR, 348606-BL) or isotype controls (VWR, 400212-BL) were added to two Eppendorf tubes each and incubated on ice for 30 minutes. Stained cells were washed twice with 1 ml of “staining buffer” and once with 1 ml PBS prior to being taken up in 100 ul of PBS. Quantitation of Pgp expression was conducted on a BD Accuri benchtop flow cytometer using phycroerythrin-conjugated calibration beads according to manufacturer’s instructions (Bangs Laboratories, #821).

### Calcein AM Uptake

DU4475 cells were plated at 100,000 cells/well in a black 96 well plate with 1 uM Calcein AM (UltraPure Grade, AnaSpec) + and – 5 uM of Pgp inhibitor Zosuquidar (SelleckChem). After addition, fluorescence (Calcein excitation = 485 nm, emission = 526 nm) was measured every 10 minutes for 3 hours at 5% CO2 and 37°C using a Synergy H1 microplate reader and CO2 controller (BioTek).

## Results

To better understand BRENDA, we extracted kcat and Km values from this database and augmented this source with manual curation of additional literature. We wanted to use BRENDA to better understand Pgp as a “vacuum cleaner” model more quantitatively (Raviv et. al., 1990). So, we searched for primary literature references with data on Km and kcat values for Pgp. This analysis produced 137 entries including 47 different compounds with publications from 1995 to 2022. The average kcat was 1.29 seconds^-1^ and Km was 154.88 uM. Our data ranged from 0.0139 to 1040 uM for the Km and 0.7 to 3.3 seconds^-1^ for the kcat. There was significantly more data on Km than kcat which reflects more scientific focus on Km as a metric for studying enzyme kinetics.

By overlaying the Pgp manual curation dataset with BRENDA for Km and kcat, the wide variability for these enzyme kinetics values is apparent. Even for the same substrate, there can be wide variety in the Km and kcat which can be traced to different experimental platforms and conditions (Fig 1). Fig 1A shows literature reported kcat and Km values for Pgp vs enzymes in BRENDA. There is no agreement in the scientific literature on a single value for these enzyme kinetics parameters although there is major overlap among the reported values.

**Fig 1.**
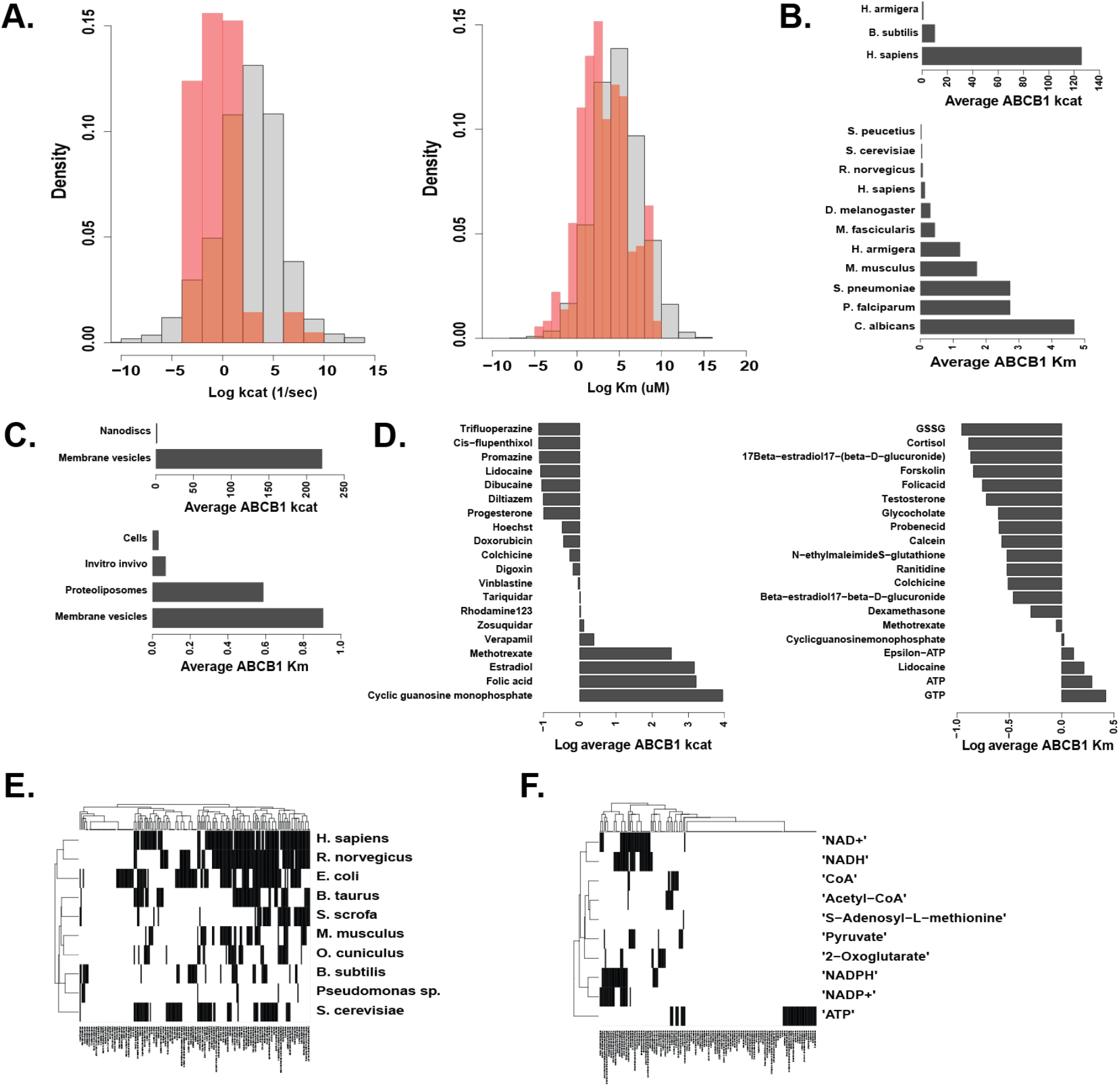
BRENDA Enzyme Kinetics Data. The BRENDA database offers enzyme information across organism, platform and compound. (A) Literature on Pgp reflects variability in reported enzyme kinetics values for kcat and Km. (B-D) Inconsistencies in enzyme information can be attributed to intrinsic differences in experimental conditions with organisms, platforms and compounds used. (E-F) BRENDA has current gaps in enzyme information across organism and compound.

Fig 1B displays the organisms within the Pgp manual curation dataset and average values for ABCB1 kcat and Km. As shown, *Homo sapiens* (humans) have highest average kcat and *Candida albicans* (yeast) have the highest average Km. Among the various experimental platforms used to study Pgp *in-vitro*, membrane vesicles yielded the highest average kcat and Km (Fig 1C). This is likely due to differences in Pgp functionality once purified and reconstituted whether in membrane vesicles, liposomes or nanodiscs. In addition, there are many choices for substrates to study Pgp including inhibitors (Verapamil, Tariquidar, Zosuquidar), fluorescent dyes (Hoechst, Calcein, Rhodamine123) and oncology drugs (Doxorubicin, Vinblastine), nucleotides (ATP, GTP) among others. Thus, the reported kcat and Km values based on compound is widespread since Pgp can bind vastly differently to a spectrum of compounds based on molecular weight, chemical structure etc. (Fig 1D).

Even though BRENDA is a comprehensive database, it still has gaps in studies. To illustrate this, the top 10 organisms (Fig 1E) and top 10 compounds (Fig 1F) in BRENDA were compared to the top 200 BRENDA enzymes by Km values. For each organism-enzyme pairing and compound-enzyme pairing, if there are any reported Km values within BRENDA, the pairing is black. But if there are no reported Km values within BRENDA, the pairing is white. As demonstrated, there are gaps in our understanding of kinetics for specific enzymes in organisms and in compounds. But the knowledge gap is much more pronounced in our understanding of enzyme kinetics regarding compounds. This indicates that variation in enzyme kinetics is more widely thought about in terms of the organism. Overall, the widespread variability in organism, experimental platform and compound contribute to the disagreement on enzyme kinetics values in the scientific literature. As a drug transporter, there are many avenues for studying Pgp functionality and substrates.

With this raw Km and kcat data in hand, we next examined the current largest database documenting MDR-substrates available. BRENDA focuses on enzymes generally, but we wanted to narrow our focus to enzymes involved in MDR. Thus, we turned to DrugBank, a database developed to bridge gaps between clinically oriented drug resources and chemically oriented drug databases. Originally, DrugBank combined clinical information (drug actions, pharmacology) with chemical information (structures, properties) in a single platform. It was tailored to pharmacologists, medicinal chemists and pharmacists. It was first released in 2006 and since then, has released five updated versions with enhanced data compilation, visualization, application etc. (Wishart et. al., 2006, Wishart et. al., 2008, Knox et. al., 2011, Law et. al., 2014, Wishart et. al., 2018, Knox et. al., 2024).

The 2011 version (DrugBank 3.0) added data on drug pathways, transporters, metabolizing enzymes, pharmacogenomics and adverse drug reactions (Knox et. al., 2011). The data on transporters and metabolizing enzymes stems from scientific literature curation which introduces experimental variability. So, for transporters and metabolizing enzymes, DrugBank defines substrates in a binary way by assigning 1’s to substrates and 0’s to non-substrates (Knox et. al., 2011). This variability is like BRENDA, so DrugBank probably used this classification method as a compromise because DrugBank curators could not find a consensus for Km or kcat values.

DrugBank has information on 63 xenobiotic enzymes. Since we are most interested in oncology drugs, we created a binary heatmap for MDR enzyme-drug pairings (Figure S4). In the binary heatmap, black indicates a substrate and white indicates a non-substrate according to DrugBank definitions. From the data, most current knowledge of MDR genes is centered on ABCB1 which is the gene for Pgp. Thus, we focused our efforts on Pgp in the broader context of understanding MDR quantitatively (https://go.drugbank.com/).

Clearly, DrugBank is a solid start to consolidating our knowledge on MDR but still has major gaps. Due to inconsistencies with MDR studies, a new metric is needed that has the potential for functionally scoring drugs by substrate specificity. Since substrate specificity is determined by gene expression and enzyme kinetics, we focused on CCLE and PRISM databases which encompass both datasets across 479 cancer lines (Ghandi et. al., 2019, Corsello et. al., 2020).

CCLE was generated from a collaboration between the Broad Institute out of MIT and Harvard and Novartis Institutes for Biomedical Research (Barretina et. al., 2012). CCLE covers the genetic characterization of ∼1,000 cancer lines and has a plethora of data types including DNA, mRNA, protein, metabolites etc. We are interested in mRNA data specifically because Pgp and other xenobiotic enzymes are regulated at the RNA level by transcription factors (Tolson et. al., 2011).

We chose to use mRNA because it is generally accepted that xenobiotic enzyme expression is regulated at the transcriptional level (Xu et. al., 2005). For example, the Pregnane X Receptor (PXR) is one transcription factor which is activated after drug binding and localized in the nucleus. PXR increases transcription of ABCB1 and is activated after initial exposure to chemotherapies indicating its role in MDR. PXR activation corresponds to Pgp protein expression and is critical to the induction of Pgp expression and efflux (Harmsen et. al., 2010, Oladimeji et. al., 2018).

Although it provides critical information, the current form of CCLE does not have much data on drugs, so we need to supplement our research with other resources. We turned to drug screening platforms and chose PRISM because of the overlap with CCLE in cell line screening. Originally, PRISM was generated from a collaboration between the Broad Institute and Sanger Institute of the United Kingdom to screen drugs across ∼500 genomically characterized cell lines. PRISM cancer lines are barcoded then pooled and drugs tested via high throughput screens that measure efficacy (Corsello et. al., 2020).

PRISM screened 1,448 compounds against 499 cell lines. We chose PRISM over other drug screening datasets such as GDSC and CTRP because PRISM had more FDA approved drugs (Corsello et. al., 2020, Yang et. al., 2013, Basu et. al., 2013). Since one of our goals is to optimize clinical drug selection, it was critical for our chosen dataset to incorporate FDA approved oncology drugs.

Combining data from CCLE and PRISM can give a functional score of MDR using strategy described in Fig S5. The Cancer Dependency Map Portal (DepMap Portal) combines CCLE and PRISM data to map genomic and genetic dependencies of drugs. DepMap Portal provides correlations between CCLE mRNA expression data and PRISM drug AUC data. Pearson coefficients from these CCLE and PRISM correlations offer a functional score of drug resistance (https://depmap.org/portal, Tsherniak et. al., 2017). Specifically, we can use our knowledge of drug pharmacodynamics (how drugs interact with cells) and Michaelis-Menten enzyme kinetics for drug transporters to mathematically model EC50 change as a linear relationship.

The assumption of 1^st^ order Michaelis-Menten conditions (consistent with clinical pharmacokinetics assumptions) make the EC50 proportional to Pgp expression. The enzyme substrate turnover rate (kcat), enzyme-substrate affinity (Km), and diffusion rate (kdiff) comprise the slope. A higher kcat means Pgp is pumping out more drug and thus increases the EC50. Similarly, a lower Km indicates greater binding affinity and increases the EC50. A higher kdiff means that more drug is getting into the cell which would decrease the EC50. As explained previously, we can approximate the EC50 to the Kd. For the MDR phenotype, a drug’s EC50 is determined by multiple MDR genes of which ABCB1 contributes. So, in a linear regression model, the slopes ^*^ expression for each drug efflux gene are additive (Fig S5A). For my research, we wanted to use ABCB1 as a proof of concept for quantifying MDR as a multigene phenotype.

The Cancer Dependency Map Portal combines CCLE mRNA expression data with PRISM drug EC50 data across ∼500 cancer cell lines. Specifically, CCLE profiles 20,000 genes and 1,000 cancer cell lines, and PRISM encompasses 1,500 drugs across 1,000 cancer cell lines (Fig S5B). Since drug EC50 correlates linearly with ABCB1 expression, the correlation coefficient (r) explains the % variance attributed to ABCB1 expression. In our analysis, each data point is a single cancer cell line (Fig S5C). In DepMap Portal, we used the Data Explorer tool to compare ABCB1 expression (log2(TPM+)) with individual oncology drug EC50 (log2 fold change) across ∼500 cancer cell lines. Under the Linear Regression section, we extracted data from the correlation including the Pearson coefficient, Spearman coefficient, Slope, Intercept and p-value for the FDA approved library of oncology drugs (Tsherniak et. al., 2017).

Fig 2A shows a general model of Michaelis-Menten enzyme kinetics for all known xenobiotic enzymes for the anthracyclines and taxanes. These xenobiotic enzymes include drug transporters and metabolism enzymes. We examined a subset of DrugBank data for the anthracyclines and taxanes (Fig 2B). We chose the anthracyclines and taxanes specifically because they are oncology drugs and frequently studied in the context of MDR. An analysis of BRENDA demonstrates no consensus Km for the parent compounds, Doxorubicin (17 uM) or Paclitaxel (0.5 uM) (Fig 2C). For each drug class, we analyzed MDR genes focusing on ABC and Solute Carrier (SLC) drug transporters and CYP450 metabolizing enzymes with drug target genes as negative controls. The anthracyclines and taxanes have different mechanisms of action and intracellular targets. The anthracyclines target topoisomerase II and reduction-oxidation enzymes because they work through DNA disruption and reactive oxygen species accumulation. The taxanes target tubulin by interfering with microtubule depolymerization during the cell cycle (DrugBank). A heatmap of binary DrugBank definitions (black = substrate, white = non-substrate) shows that most knowledge on the anthracyclines and taxanes is concentrated on the parent compounds, Doxorubicin and Paclitaxel (Fig 2D).

**Fig 2.**
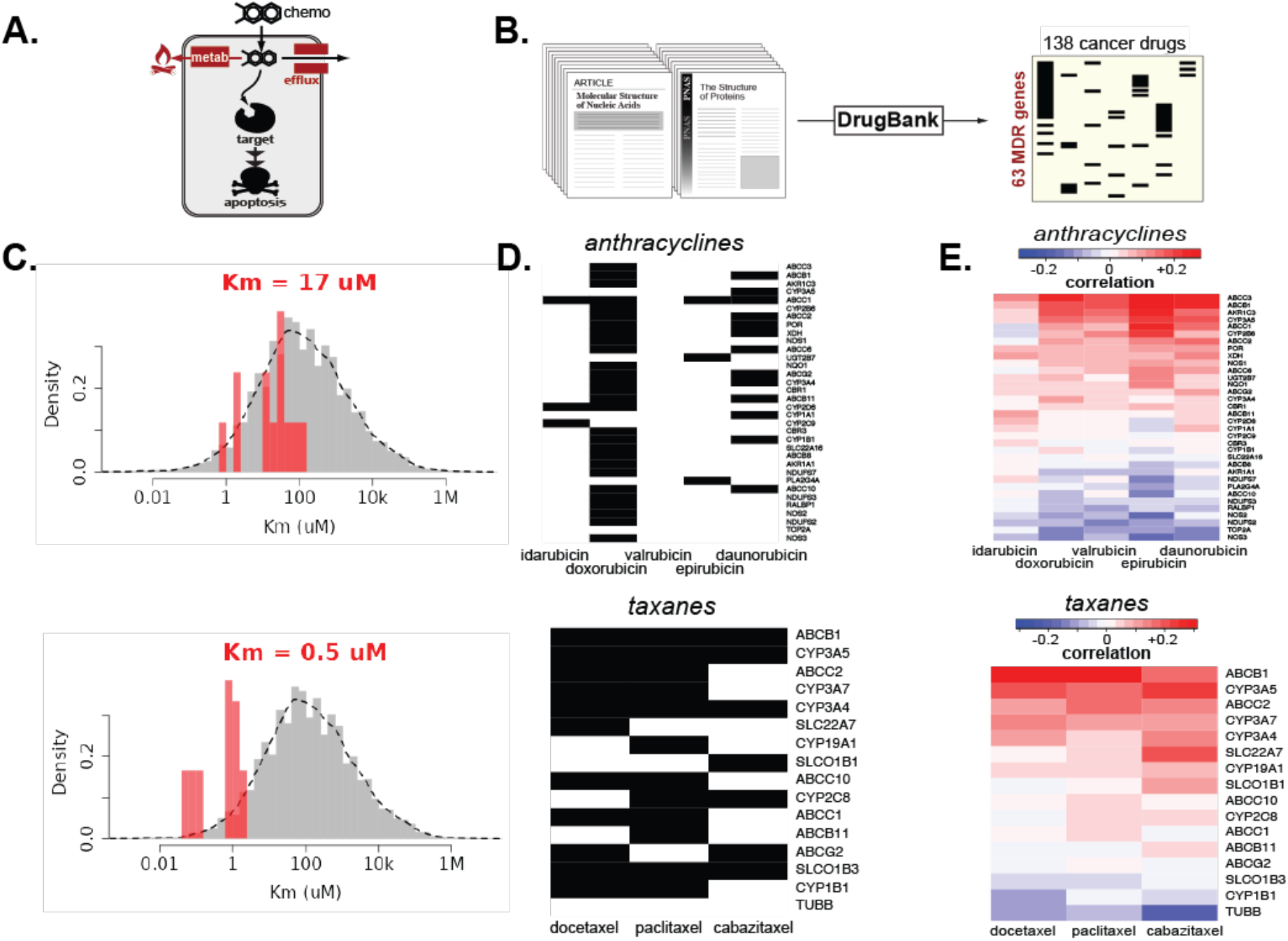
Correlation Analysis to Fill Gaps. Our correlation analysis fills in literature gaps and inconsistencies. (A) A fraction of drug causes cytotoxicity after efflux and metabolism. (B) DrugBank literature assigns binary definitions to Pgp substrates. (C) Literature values for Km of Doxorubicin (17 uM) and Paclitaxel (0.5 uM) are inconsistent. (D) MDR gene-drug pairs for the anthracyclines and taxanes using DrugBank definitions. (E) MDR gene-drug pairs for the anthracyclines and taxanes using our correlation analysis.

But many gaps exist in understanding most oncology drugs and MDR genes. A pilot study of our correlation analysis fills in these gaps and gives a quantitative ranking of xenobiotic enzymes with ABCB1 at the top where a higher ranking indicates greater substrate specificity (Fig 2E). Our correlation analysis fills those gaps by bridging gene expression and drug EC50 through CCLE and PRISM datasets. Regenerating functional heatmaps for the same MDR genes and drugs gives a continuum of substrate specificity. For the anthracyclines and taxanes, the top three genes with the highest Pearson coefficient include ABCB1. Therefore, we focus on ABCB1 because of its wide application in understanding MDR and oncology drugs.

To investigate the function of Pgp, we analyzed 20 oncology drugs in the FDA approved library across 479 cancer cell lines. These cell lines were chosen because they overlapped in CCLE and PRISM datasets. Both CCLE and PRISM datasets were downloaded and analyzed using the R-statistical language and new matrices created with the 500 overlapping lines for ABCB1 and each drug. Linear regression analysis compared CCLE ABCB1 expression with PRISM drug EC50 for the top 20 oncology drugs. Most drugs had a positive correlation with ABCB1 which is consistent with Pgp as an enzyme that causes some resistance.

We expanded this analysis to all 138 oncology drugs in the FDA approved library and to other ABC genes such as ABCC2 and ABCG2. Since MDR is a multigene phenotype, we expect different drugs to be resisted by different genes (Fig S6A). Each MDR gene acts like “armor” to shield the cancer cell from oncology drugs. However, each MDR “armor” is best equipped to “shield” cancer cells from a subset of drugs. Our quantitative rankings help identify which oncology drugs are most and least likely to pierce the MDR “armor worn “by cancer cells. Our analysis generated waterfall plots which rank 138 FDA approved oncology drugs by their Pearson coefficients which were determined from correlation analyses between ABC gene expression and drug EC50 across 479 cancer cell lines. Thus, the Pearson coefficient is a measure of substrate specificity for each drug-gene pair. A greater Pearson coefficient indicates greater substrate specificity and a higher probability of drug resistance.

Since our focus is on taxanes, vinca alkaloids and anthracyclines, we highlighted these drugs in the waterfall plots (Fig S6B). For ABCB1 and ABCC2, most of these drugs are highly ranked except for Idarubicin which may be less of a Pgp substrate than other anthracycline drugs (Roovers et. al., 1999, Smeets et. al., 1999). Though similar, there is some variation in the rankings between ABCB1 and ABCC2 which represents differences in preferred substrates for each enzyme. Many factors contribute to substrate preferences such as the drug molecular weight and chemical structure, binding affinity and enzyme binding pocket. The enzyme for ABCG2, Breast Cancer Resistant Protein (BCRP), is much different than Pgp which is reflected in its quantitative rankings of oncology drugs (Robey et. al., 2018). ABCG2 has a much narrower binding pocket, so drugs which are strong substrates of ABCB1 are only moderate or weak substrates of ABCG2. With this analysis between gene expression and drug potency, EC50 is not always measurable, so we used drug area under the curve (AUC) as a surrogate which is more common (Corsello et. al., 2020, Yang et. al., 2013, Basu et. al., 2013). The AUC has limits which are described more in Fig S8.

Since most drugs had a positive correlation with ABCB1 expression, we wanted to determine if quantitatively defining Pgp substrates agreed with the scientific literature. So, we used CCLE and PRISM which has Pgp expression and drug AUC data across 479 cancer lines (Fig 3). Then, we separated 138 drugs which were listed in DrugBank and the FDA approved oncology library based on their binary definitions. Once we grouped the drugs into ‘Pgp Known’ (70 drugs) or ‘Pgp Undefined’ (68 drugs) by DrugBank definitions (Knox et. al., 2011), we input their Pearson coefficients from correlations between ABCB1 expression and drug AUC. Then, we took the average Pearson coefficient from each group and conducted a one-tailed Student’s T-test to determine if DrugBank binary definitions aligned with quantitative Pearson coefficients (Fig 3D). The T-test confirmed that Pearson coefficients correctly identified Pgp substrates in agreement with DrugBank (p-value = 0.0129). Critically, the Pgp Known drugs had a higher average Pearson coefficient than the Pgp Undefined drugs (Fig 3D).

**Fig 3.**
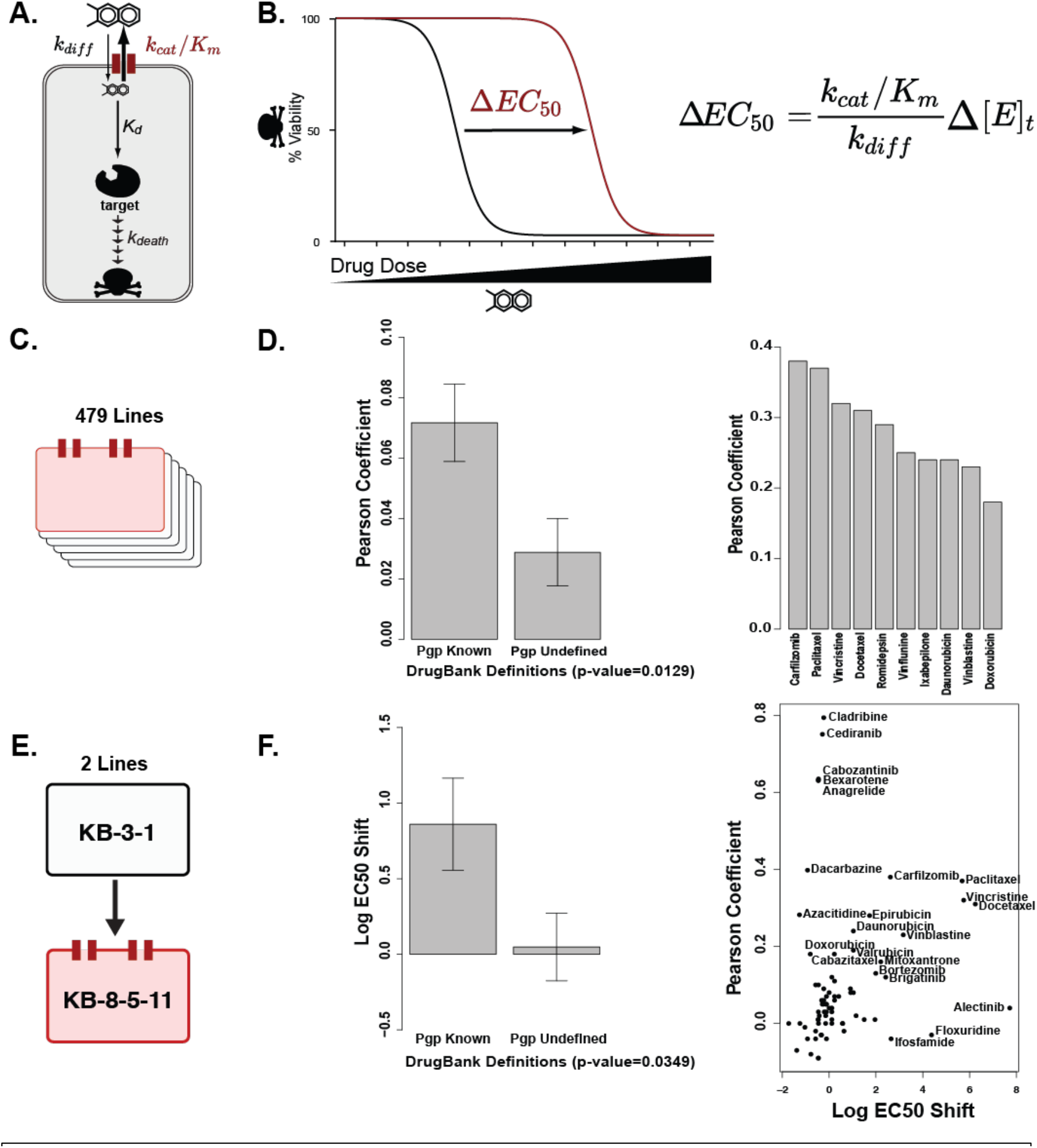
Quantitative Metrics for Substrate Specificity. (A) MDR is modeled through diffusion rate (kdiff), enzyme kinetics (kcat/Km) and MDR expression. (B) Increased MDR expression shifts the EC50 right. (C) Our correlation analysis covers 479 cancer lines with natural Pgp expression. (D) DrugBank and Pearson coefficients agree on Pgp substrate definitions. The top 10 ranked oncology drugs by Pearson coefficient include taxanes, vinca alkaloids and anthracyclines. (E) Lee *et. al*. conducted a drug screen of 10,804 compounds using two cervical cancer lines with induced Pgp expression. (F) DrugBank and log EC50 shift agree on Pgp substrate definitions.

We then ranked the FDA approved library of 138 oncology drugs in descending order by their Pearson coefficients. Since Pearson coefficients are a measure of Pgp substrate specificity, the drugs to which Pgp confers the most resistance should be the highest ranked. Our general analysis yielded >85% drugs with a positive correlation. We focused on the top 10 oncology drugs which are enriched in the taxanes (Paclitaxel, Docetaxel), vinca alkaloids (Vincristine, Vinflunine, Vinblastine) and anthracyclines (Daunorubicin, Doxorubicin). The other top 10 drugs were Carfilzomib, a proteasome inhibitor, Romidepsin, a histone deacetylase inhibitor and Ixabepilone, an epothilone (Fig 3D). There is some variety in the top 10 drugs in class and mechanism of action supporting Pgp conferring resistance to several oncology drugs.

Our computational analysis linked ABCB1 expression to drug AUC, but not all cell lines and drugs yielded data. In the PRISM drug screen, some drugs were not cytotoxic. Therefore, we wanted to look at EC50 specifically as a quantitative metric. Additionally, we wanted to study how changes in Pgp expression changes the drug EC50. Compared to our approach, Lee *et. al*. utilized two cell lines with induced Pgp expression which simplifies the study of Pgp expression on drug potency but might not accurately represent a model of acquired multidrug resistance through increased Pgp expression (Lee et. al., 2019).

In this study, a high throughput screen (HTS) was conducted using compound libraries to determine cytotoxicity with differential Pgp expression (Fig 3E and 3F). These researchers used a parent KB-3-1 human cervical adenocarcinoma and Pgp overexpression subline KB-8-5-11 which they induced through incremental dosing with Colchicine (100 ng/ml) and verified through flow cytometry. For the HTS, compounds were screened against the parent line and Pgp overexpression line – and + Pgp inhibitor Tariquidar to measure the reversibility of drug resistance to compounds which were classified as Pgp drug substrates (Lee et. al., 2019).

Compounds were taken from the Mechanism Interrogation Plate (MIPE) library (1,912), National Center for Advancing Translational Science pharmaceutical collection (NPC) (2,816), NCATS Pharmacologically Active Chemical Toolbox (NPACT) (5,099) and kinase inhibitor library (977). The MIPE library includes oncology compounds at preclinical, investigational and FDA approved stages. The NPC library contains compounds approved by the FDA and some drugs approved by agencies in other countries. The NPACT library has compounds with an emphasis on novel phenotypes, biological pathways and cellular processes (ncats.nih.gov/research/research-activities/compound-management).

For the HTS, cells were plated at 500 cells/well in 1536 well plates in 5 ul of media and incubated at 37 °C and 5% CO2 with compound for 72 hours. After 72 hours, Cell Titer-Glo reagent was added to all wells, incubated for 5 minutes and luminescence read. The drug screen yielded 90 of 10,804 compounds which were Pgp substrates and 55 of the 90 Pgp substrates which were novel. For the KB-3-1 parent line, 1,362 of the 10,804 compounds were cytotoxic. Thirty percent of the kinase, 21% of the MIPE, 10% of the NPACT and 5% of the NPC compound libraries exhibited cytotoxicity in the HTS (Lee et. al., 2019).

Lee *et. al*. induced Pgp to study the effect of expression on drug EC50 or [drug] required to kill 50% of cells which is a measure of cytotoxicity. The difference between dose response curves in the parent KB-3-1 and Pgp overexpression KB-8-5-11 lines is the AUC. The ΔAUC is correlated to the ΔPgp expression and drug efflux between the cell lines. In the HTS, they gathered dose response data for the KB-3-1 parent line, KB-8-5-11 Pgp overexpression line and KB-8-5-11 line + 1 uM Tariquidar to measure reversibility of drug resistance.

Dose response curves were plotted by the log drug (M) on the x-axis and % Activity on the y-axis. The % Activity corresponds to cell viability where -50 means 50% cell death. Additionally, they measured the ΔAUC between the KB-3-1 parent line and KB-8-5-11 Pgp overexpression line as ΔAUC1. The ΔAUC between the KB-8-5-11 line and + 1 uM Tariquidar is ΔAUC2. For the 90 compounds identified as Pgp substrates, a rank order analysis was conducted which plotted ΔAUC1 against ΔAUC2 for the 90 Pgp substrates with a positive correlation (R^2^ = 0.69) (Lee et. al., 2019).

For the HTS, our lab created an online app to visualize dose response curve data in KB-3-1 and KB-8-5-11 lines for 10,804 compounds. Our online app calculates the ΔEC50 (fold change) in the drug-resistant vs parent lines. A greater fold change indicates stronger Pgp substrate specificity (https://douglasslab.shinyapps.io/mdr_screen/).

Since the data from Lee *et. al*. links drug ΔEC50 to Pgp expression, we wanted to determine if ΔEC50 agreed with Pgp substrates as defined by the scientific literature. So, we grouped the FDA approved oncology drug library into ‘Pgp Known’ if DrugBank classified them as Pgp substrates (70 drugs) and ‘Pgp Undefined’ if DrugBank classified them as Pgp non-substrates (68 drugs). Then, we listed the log ΔEC50 from Lee *et. al*. for the 138 oncology drugs and calculated the average log ΔEC50 for each group.

We conducted a one-tailed Student’s T-test comparing DrugBank vs log ΔEC50 substrate definitions for each group. The T-test yielded a p-value of 0.0349 which demonstrated that log ΔEC50 can be used as a quantitative metric for assessing Pgp substrate specificity. As with the Pearson coefficient, the Pgp Known drugs had a higher log ΔEC50 than the Pgp Undefined drugs. As quantitative metrics, both Pearson coefficients and log ΔEC50 incorporate Pgp expression which is critical to assessing MDR. Since tissues have differential expression of MDR genes, incorporating expression into quantitative metrics of substrate specificity is more accurate and predictive of drug resistance.

Although both metrics incorporate Pgp expression, the difference is Pearson coefficients come from natural Pgp expression vs log ΔEC50 coming from induced Pgp expression. Additionally, Pearson coefficients were generated from an analysis of 479 cancer lines vs log ΔEC50 from an analysis of two cancer lines (Fig S7A and S7C). So, we then wanted to assess agreement between Pearson coefficients and log ΔEC50 as quantitative metrics of Pgp substrate specificity. We generated a scatter plot to study the correlation between Pearson coefficients and log ΔEC50 with 138 FDA approved oncology drugs.

For the scatter plot, 138 drugs were plotted by their log ΔEC50 on the x-axis and Pearson coefficient on the y-axis. Our analysis includes a cluster of data points (drugs) with low log ΔEC50 and Pearson coefficients indicating these are likely weak or non-Pgp substrates. The rest of the drugs had higher log ΔEC50 and Pearson coefficients meaning they are likely stronger or moderate Pgp substrates. In general, there is a positive correlation between both quantitative metrics. Some drugs exhibit stronger substrate specificity as measured by log ΔEC50 or Pearson coefficient which could be due to differences in cell lines. Lee *et. al*. used two ovarian cancer lines for their HTS vs 479 cancer lines for the PRISM screen. This is evidence supporting the use of multiple quantitative metrics for the most comprehensive understanding of MDR.

Even though the PRISM screen covers more cell lines, obtaining dose response data only works when the drug is cytotoxic. If the drug is not cytotoxic, there will not be a dose response curve or EC50. Once a drug enters a cell, it must bind to the intracellular target and cause cell death within the range of drug concentrations tested (Fig S8A).

Earlier, we conducted a linear regression analysis of the top 20 oncology drugs comparing CCLE ABCB1 expression on the x-axis with drug AUC on the y-axis. We can use Paclitaxel as an example of the limitations of the PRISM screen in some cell lines. In the linear regression analysis, there are vertical (black box) and horizontal (red box) clusters of cancer lines. In the vertical cluster, these cell lines have low to no ABCB1 expression and a range of AUCs for Paclitaxel. These cell lines likely have higher drug sensitivity to Paclitaxel. In the horizontal cluster, these cell lines have low to high ABCB1 expression and an AUC = 1 for Paclitaxel. Paclitaxel resistance in the low ABCB1 expression cell lines is not due to ABCB1 but other drug transporters or metabolizing enzymes. Paclitaxel resistance in the high ABCB1 expression cell lines is due to ABCB1 (Fig S8B).

As shown, AUC is a surrogate measurement of drug EC50. For an AUC = 1, there is no dose response curve within the drug concentration range. This phenomenon can represent a few different scenarios. The first scenario is that a higher drug concentration is needed for cytotoxicity which is outside the drug screen concentrations as shown by the dotted line extrapolation (Fig S8C). In drug screens, a standard range is typically used, and less potent drugs require a higher concentration to be cytotoxic. The second scenario is that a drug is non-toxic to the cell line regardless of concentration. This would be a likely explanation for drugs whose mechanisms of action do not cause cytotoxicity. However, we limited our analysis to oncology drugs which are cytotoxic, so the first scenario is the most plausible.

Since PRISM has potential false negatives because of dose range limitations, we investigated studies which use fluorescent probes and Pgp inhibitors to assess substrate specificity. We found a study which looked at MDR in multiple myeloma with anthracycline drugs and a Pgp modulator. Anthracyclines are oncology drugs and of great interest to us given our computational analysis classifying them as strong Pgp substrates. Thus, this paper is particularly relevant for a direct comparison with our results. Roovers *et. al*. studied a newer anthracycline, Idarubicin, in the context of multiple myeloma to see if its higher lipophilicity could increase cytotoxicity compared to Doxorubicin and Daunorubicin (Roovers, et. al., 1999). With a higher lipophilicity, Idarubicin has the potential for better efficacy and potency in multiple myeloma treatment.

A parent multiple myeloma cell line 8226-S and two Pgp overexpression cell lines were chosen to assess cytotoxicity and uptake kinetics and Doxorubicin, Daunorubicin and Idarubicin. Pgp overexpression was induced in 8226-R7 cells with a single dose of Daunorubicin and 8226-Dox40 cells with multiple doses of Doxorubicin. This strategy is analogous to the strategy used in Lee *et. al*. to induce Pgp expression in the Pgp overexpression cancer lines (Roovers et. al., 1999, Lee et. al., 2019).

Dose response and cytotoxicity were assessed after a 3-day incubation + drug + or - Pgp modulator, Verapamil, through the MTT colorimetric assay. The MTT assay is based on the reduction of the yellow tetrazolium salt, 3-(4,5-dimethylthiazol-2-yl)-2,5-diphenyltetrazolium bromide (MTT), to a purple formazan crystal by live cells. Optical density (OD) was measured at 750 nm where a higher OD corresponds to more live cells (less transparent) and a lower OD indicates more cell death (more transparent). Dose response curves were plotted as drug concentration (uM) on the x-axis and OD750 on the y-axis. Data was gathered for Doxorubicin, Daunorubicin and Idarubicin across the 3 cell lines (Roovers et. al., 1999).

The 8226-S parent line exhibited the greatest cytotoxicity with Idarubicin having the most potency and Doxorubicin having the least potency. The 8226-R7 Pgp overexpression line demonstrated more resistance to all three drugs, decreasing their potency (Roovers et. al., 1999). Idarubicin still had the best potency and Doxorubicin the least potency. Lastly, the 8226-Dox40 Pgp overexpression line had even more resistance to all three drugs, with no drug achieving 100% cell death at the highest drug concentration (4 uM) (Roovers et. al., 1999). However, Idarubicin still had the greatest potency of the three drugs. Since drug potency decreased with increasing Pgp expression, it follows that Idarubicin could be a weak Pgp substrate. The ΔEC50 for Idarubicin was much less than for Doxorubicin and Daunorubicin across the three cell lines (Roovers et. al., 1999).

The same experiment was conducted with the addition of 50 uM Verapamil, a 1^st^ generation Pgp inhibitor. In theory, the stronger Pgp substrates, Doxorubicin and Daunorubicin, will exhibit more potency with Verapamil in the Pgp overexpression cell lines. As expected, the addition of Verapamil did not significantly change the dose response curves in the 8226-S parent line. In the 8226-R7 line, Doxorubicin and Daunorubicin were slightly more potent than without Verapamil. In the 8226-Dox40 line, the difference was more pronounced where the addition of Verapamil achieved 100% cell death with Doxorubicin, Daunorubicin and Idarubicin at 2 uM and 4 uM of drug (Roovers et. al., 1999). These results emphasize optimal drug selection for cancer treatment. Pgp inhibitors such as Verapamil are best used in Pgp-high cell lines, and drugs which are resisted by Pgp need to be given in cancers with low or no Pgp for optimal clinical results.

To measure Pgp expression, Roovers *et. al*. used the anthracyclines as fluorescent probes and flow cytometry across 8226-S, 8226-R7 and 8226-Dox40 lines (Fig 4A). Essentially, the anthracyclines’ fluorescence is an indirect measure of drug accumulation inside cells. As Pgp expression increases, more anthracycline can be pumped out and the fluorescence decreases. So, the Δ[E] is directly proportional to the Δ[Probe].

**Fig 4.**
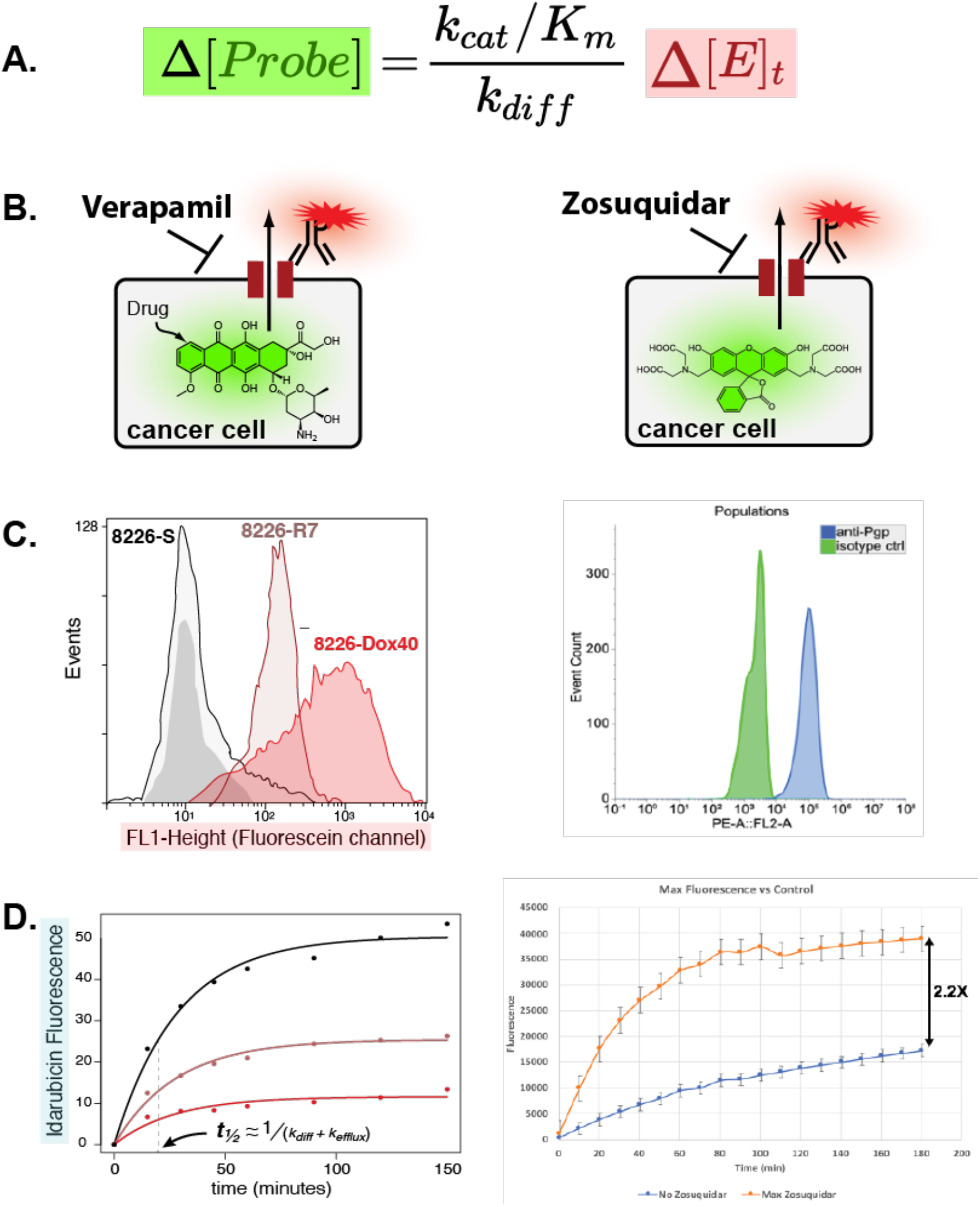
Fluorescent Probes for Drug Screening. (A) Probe fluorescence inversely changes with MDR expression where lower fluorescence indicates higher expression. (B) Pgp substrate specificity is measured through intracellular accumulation of drug (Roovers *et. al*.) or fluorescent dye (Ashley) +/-addition of Pgp inhibitor. (C) Flow cytometry quantifies Pgp expression in multiple myeloma (Roovers *et. al*.) or breast cancer (Ashley) cell lines through antibody staining. (D) Fluorescent microscopy tracks uptake of Idarubicin (Roovers *et. al*.) or Zosuquidar (Ashley) into cells over time. The kinetics of uptake are modeled through k_diff_ and k_efflux_.

They used flow cytometry to quantify Pgp protein expression across all three multiple myeloma cell lines (Fig 4C). For this analysis, 4 ^*^ 10^5^ cells were incubated with 10 ul of 50 ug/ml mouse Pgp monoclonal antibody MRK16 for 60 minutes at room temperature. Cells were then washed with a phosphate buffered saline-fetal calf serum (PBS-FCS) buffer and incubated for 15 minutes at room temperature with 0.5 ug of goat anti-mouse IgG2a-FITC. Results confirmed the highest Pgp expression in 8226-Dox40 cells followed by slightly less expression in 8226-R7 cells and basal expression in the 8226-S cells (Roovers et. al., 1999). Since flow cytometry measures Pgp expression, it confirms the quantify of Pgp protein at the cell surface but does not assess functionality of the enzyme as a drug transporter.

To determine Pgp functionality, Rhodamine123 was used as a fluorescent probe to measure efflux. Rhodamine123 is a cell permeable, green, fluorescent dye that is also a known Pgp substrate. In these studies, the Rhodamine123 efflux ratio (Rho123 fluorescence – Verapamil / Rho123 fluorescence + Verapamil) was determined where a smaller efflux ratio indicates greater Pgp functionality (Fig 4B). The cells were washed, resuspended in culture medium at 8 ^*^ 10^5^ cells/ml with 125 nM Rh123 and incubated at 37 °C for 10 minutes. The cells were washed twice, resuspended in culture medium and plated in 24 well plates at 0.5 cells/well. After a 20-minute CO2 incubation, the cells were pelleted, washed and resuspended in 500 ul of PBS-FCS and run through flow cytometry (Roovers et. al., 1999). Using 8226-S cells as a control (efflux ratio = 1), the Pgp overexpression lines had lower efflux ratios (30% in 8226-R7, 10% in 8226-Dox40) which corresponded with Pgp function.

Given its higher lipophilicity and better potency, uptake kinetics were measured for Daunorubicin and Idarubicin. Chemically, Daunorubicin is more similar in structure to Idarubicin than Doxorubicin, so Daunorubicin was included for comparison. In this experiment, 1 uM of anthracycline was added to 8226-S, 8226-R7 and 8226-Dox40 cells and drug uptake monitored with fluorescent microscopy. Drug uptake measurements were taken periodically over 120 minutes. Data was plotted as time (minutes) on the x-axis vs mean fluorescence intensity (MFI) on the y-axis (Roovers et. al., 1999). After 60 minutes, the MFI had peaked in the three cell lines indicating the maximum rate of drug uptake (Fig 4D).

This curve can be used to calculate the t1/2 or time taken for half-maximal drug accumulation. Since the t1/2 is determined by both diffusion and efflux, it can be represented as 1 / (kdiff + kefflux) where greater rates indicate less time for maximum accumulation of drug. However, if little to no drug is accumulated as with Pgp-high cells, the maximum fluorescence will be relatively low. As expected, the 8226-S parent line had the greatest fluorescence indicating the most intracellular accumulation. The 8226-R7 cells had the next greatest fluorescence followed by the 8226-Dox40 cells. Relative to Daunorubicin, Idarubicin accumulation occurred seven times faster which could be attributed to Idarubicin’s higher lipophilicity (Roovers et. al., 1999).

For Roovers *et. al*., they studied intracellular concentration and MDR at the same time because the anthracyclines are naturally fluorescent Pgp substrates. But the MTT assay does not work for non-toxic compounds, and most oncology drugs are not naturally fluorescent. So, we wanted to develop a competitive assay with a known fluorophore and Pgp substrate to estimate intracellular concentrations of drug for non-toxic and non-fluorescent compounds.

With the AUC limitation of PRISM, we chose Calcein acetomethylester (AM) as a fluorescent probe to assess Pgp substrate specificity *in-vitro*. Calcein AM is a known Pgp substrate, cell permeable, fluorescent green dye (Eneroth et. al., 2001). Calcein AM easily diffuses into cells where the AM is cleaved by esterase enzymes in live cells. Once the AM is cleaved, Calcein fluoresces green inside cells. Without the acetomethylester component, Calcein cannot leave the cell without being pumped out by Pgp (Eneroth et. al., 2001).

For our work, we chose DU4475 as a Pgp-high cancer line. DU4475 is a ductal adenocarcinoma with the highest Pgp expression of 71 breast cancer lines and fourth highest Pgp expression of 1,000 cancer lines. DU4475 was used as our cell line model because of its high Pgp expression and low expression of other ABC genes including ABCC1 and ABCG2 which could be confounding variables (depmap.org/portal/, Tsherniak et. al., 2017).

We chose drugs for the screen based on Pgp substrate specificity rankings from their Pearson coefficients, so the screen included 76 strong, moderate, weak and non-Pgp substrates. The premise of the screen is a competitive binding assay where Calcein competes with drug for binding to Pgp. The intracellular accumulation of Calcein and resulting fluorescence is a measure of Pgp substrate specificity. The higher fluorescence will be achieved with drugs that are stronger Pgp substrates as they will outcompete Calcein for binding to Pgp. We began this work with fluorescent microscopy to optimize cell culture conditions and Calcein concentration, then transitioned to a 96 well plate assay and converted it into a drug screen (Fig S9).

For the inhibitor screen, 1^st^ to 3^rd^ generation Pgp inhibitors were used as positive controls for Calcein accumulation and fluorescence. These included 1^st^ generation Pgp inhibitors Cyclosporine, Nicardipine and Verapamil and 3^rd^ generation Pgp inhibitors Dofequidar, Elacridar, Encequidar, ONT-093, PGP-4008, Tariquidar and Zosuquidar (SelleckChem). Results showed that the best signal to noise was achieved with 5 uM Pgp inhibitor. As expected in a Pgp-high line, the 3^rd^ generation Pgp inhibitors demonstrated the best efficacy measured as maximum fluorescence with Dofequidar demonstrating the best potency (Fig 5).

**Fig 5.**
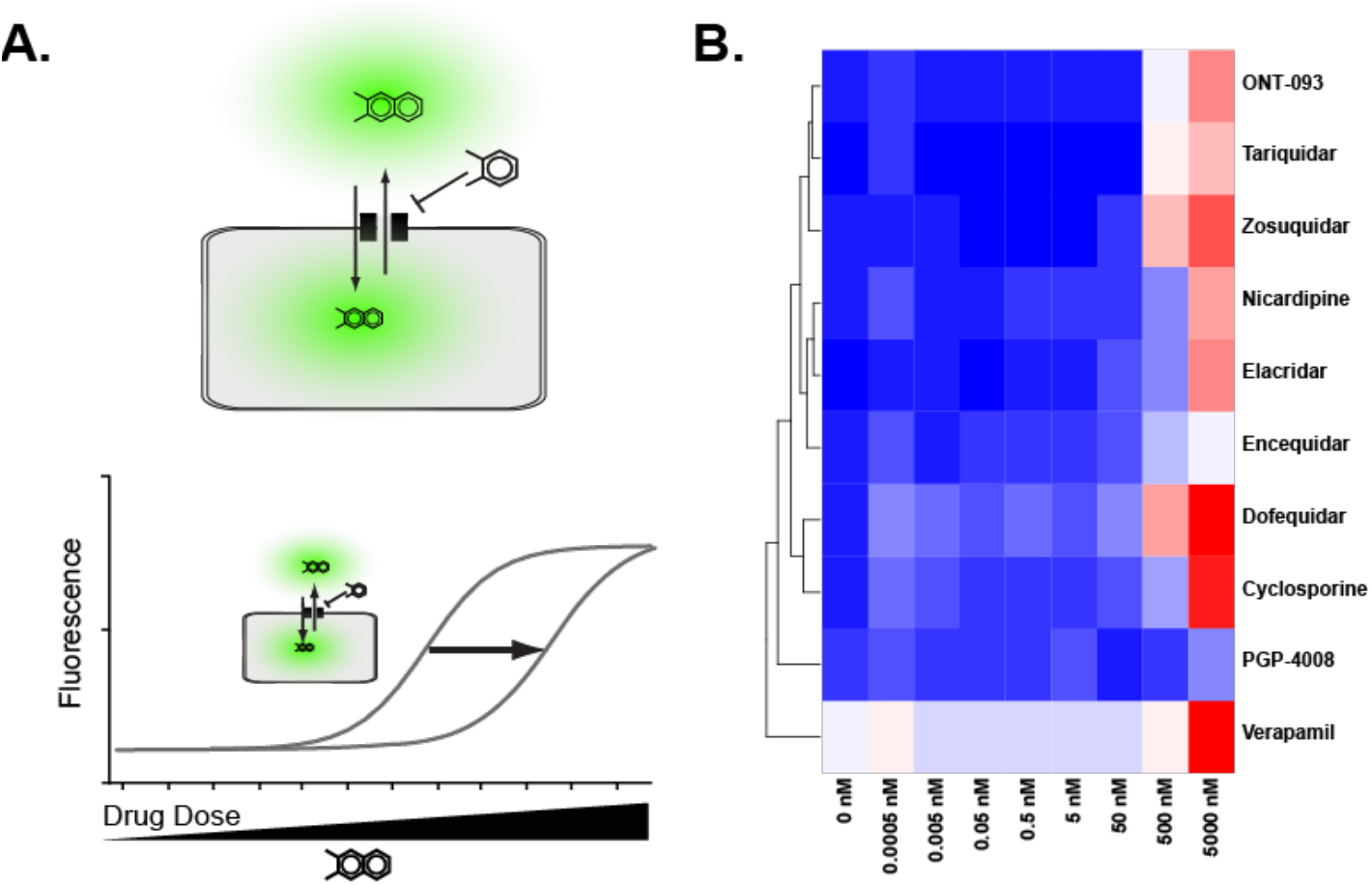
Drug Screen Measures Potency. Our Calcein AM plate-based screen measures drug potency. (A) The assay works by measuring substrate specificity through Pgp competitive binding between Calcein and drug. Potency is assessed by the drug dose at which fluorescence (Calcein intracellular accumulation) is half-maximal. (B) Our drug screen measured fluorescence dose response for 10 1^st^ and 3^rd^ generation Pgp inhibitors from 0.0005 to 5000 nM.

For the drug screen, a range of oncology drugs (SelleckChem) were chosen incorporating strong, moderate, weak and non-Pgp substrates. A total of 76 oncology drugs were screened in a competitive binding assay with Calcein AM. Results identified many top Pgp substrates in agreement with our correlation analysis such as Carfilzomib and Vinblastine. It also identified some kinase inhibitors such as Cobimetinib and Tivozanib as Pgp substrates (Fig 6).

**Fig 6.**
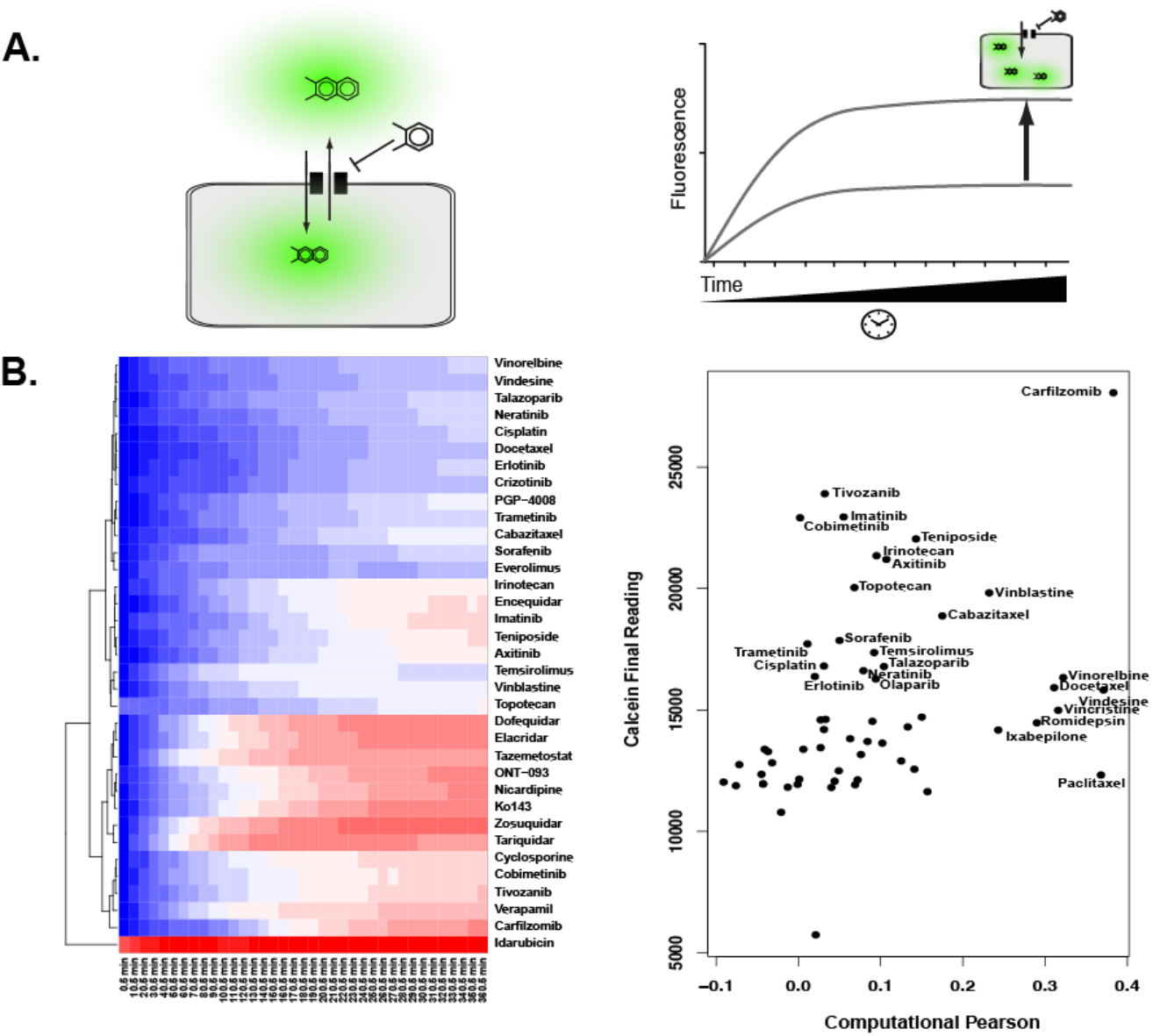
Drug Screen Measures Efficacy. Our Calcein AM plate-based screen measures drug efficacy. (A) The assay works by measuring substrate specificity through Pgp competitive binding between Calcein and drug. Efficacy is assessed by the maximum fluorescence (Calcein intracellular accumulation, 100% competition). (B) Our drug screen measured fluorescence for 76 oncology drugs across 360.5 min with readings taken every 10 min. Pearson coefficients positively correlate with drug efficacy from the Calcein assay for strong Pgp substrates.

Using beads and ∼10,000 single cells, flow cytometry analysis quantified ∼50,000 Pgp per cell in the DU4475 line. Results of Pgp isotype control and Pgp-stained with phycoerythrin antibody indicates two distinct populations which verifies a higher expression of Pgp in DU4475 cells (Fig 4C).

Calcein AM uptake was assessed using Pgp inhibitor Zosuquidar with data plotted as time (minutes) on the x-axis vs fluorescence on the y-axis. Zosuquidar was chosen as a selective Pgp inhibitor because it has been well characterized in the scientific literature (Fig 4B). At endpoint, there was a 2.2X difference in Calcein uptake within DU4475 cells based on fluorescence + and - Zosuquidar. Additionally, this analysis measures the time needed for uptake to occur which was ∼60 minutes for DU4475 cells + Zosuquidar (Fig 4D).

This analysis mirrors Roovers *et. al*. data for the 8226-S and 8226-Dox40 cell lines where the higher fluorescence in the 8226-S parent line indicates lower rates of diffusion and efflux from less Pgp expression. In our study, the higher fluorescence in the + Zosuquidar group indicates lower rates of diffusion and efflux from selective Pgp inhibition.

## Conclusions

Lee *et. al*. and our analysis both used cell-based quantitative metrics (log EC50 shift or Pearson coefficient) to assess Pgp substrate specificity. Both quantitative metrics statistically agree with DrugBank binary definitions of Pgp substrates. Additionally, the measurability of these quantitative metrics is dependent on the cellular phenotype which makes these more likely to be physiologically relevant and coupled to the phenotype of drug resistance.

In contrast, Lee *et. al*. induced Pgp expression vs our use of cancer lines with natural Pgp expression. Lee *et. al*. used two cell lines whereas we used a combination of databases which covered 479 cell lines. Lee *et. al*. investigated oncology and non-oncology drugs in their 10,804-compound screen. We focused exclusively on oncology drugs in our 76-drug screening assay.

Natural Pgp expression more accurately reflects actual cellular physiology with multiple drug transporters. But for studying Pgp, two cell lines is easier to manage and directly compare Pgp expression. For assessing MDR more generally, 479 cell lines offers a more comprehensive analysis and is widely applicable to multiple cancers. For the drug screen, including non-oncology drugs allowed for the identification of novel Pgp substrates which is critical to extending our understanding of Pgp as a drug pump.

Roovers *et. al*. and our analysis used flow cytometry to quantify Pgp expression in all cell lines using antibody. Additionally, both studies used Pgp inhibitors to assess the functionality of Pgp protein at the cell membrane surface. Both measured uptake kinetics of fluorescent drug or Calcein as fluorescent probes over time.

In contrast, Roovers *et. al*. induced Pgp expression vs our analysis which used a cell line with naturally high Pgp expression. Although both studies used Pgp inhibitors, Roovers *et. al*. used the 1^st^ generation Pgp modulator, Verapamil, and we used a 3^rd^ generation selective Pgp inhibitor, Zosuquidar. Roovers *et. al*. obtained three cell lines of various Pgp expression to study efflux. Alternatively, we used one cell line of high Pgp expression. Since their work focused on the anthracyclines, Roovers *et. al*. could use these drugs as fluorescent probes. Most of our work includes non-fluorescent drugs, so we used Calcein AM as a surrogate fluorescent probe.

Natural Pgp expression more accurately reflects cellular physiology but could introduce complexities for studying Pgp. As a selective Pgp inhibitor, Zosuquidar is better for assessing Pgp specifically whereas Verapamil is better for assessing MDR from multiple drug pumps as a Pgp modulator. Having three cell lines of incrementally higher Pgp expression is better correlated to dose response curves than a single cell line. Lastly, using drug as a fluorescent probe is simpler, but Calcein AM allows for the study of non-fluorescent drugs which comprise most of the FDA approved library.

To conclude, we can revisit the metrics presented in this work by reconciling computational and experimental definitions to more comprehensively define MDR. As a database, DrugBank is naturally tailored to clinicians with information but has expanded to include more data on pharmacokinetics and enzyme substrate specificity (Knox et. al., 2011). Unfortunately, due to its reliance on the scientific literature, DrugBank is limited to binary definitions of MDR substrates to create a consistent metric across experimental platforms.

CCLE offers mRNA expression data on ∼1,000 cancer lines. PRISM offers drug screening data on ∼500 of the CCLE cancer lines through the DepMap Portal web interface (Ghandi et. al., 2019, Corsello et. al., 2020). Combining CCLE and PRISM data offers a linear correlation between gene expression and drug AUC within the context of MDR. This linear correlation yields Pearson coefficients which are a functional, cell-based metric of enzyme substrate specificity. Pearson coefficients enable continuous drug classification where larger coefficients indicate stronger substrate specificity for MDR enzymes.

As a cancer phenotype, MDR depends on multiple factors including gene expression, enzyme kinetics and diffusion. For each cell and tissue, these factors have wide variability due to natural physiological roles, cellular phenotypes and function. These databases help reconcile these differences and contribute to a more complete picture of MDR generally. Bridging gene expression with enzyme kinetics is critical to redefining MDR quantitatively for optimizing drug selection in the clinic.

Prior scientific literature provided the foundation for EC50 (potency) and fluorescence (efficacy) as experimental metrics for assessing MDR substrate specificity. A combination of *in-vitro* approaches including flow cytometry, fluorescent microscopy and drug screens is necessary for studying Pgp substrate specificity. Pgp substrate specificity cannot be quantified by a single metric, but our analysis incorporates diffusion and enzyme kinetics into our EC50 experimental metric. By taking drug EC50 and dividing it by gene expression, we can standardize drug EC50 to any tissue or cancer.

The current research provides the foundation for measuring MDR quantitatively as a proof-of-concept through Pgp. In the future, this work can be expanded to elucidate the quantitative contributions of other non-ABCB1 ABC genes to the MDR phenotype in cancer.

## Acknowledgements

The authors would like to thank the University of Georgia Department of Pharmaceutical and Biomedical Sciences for the funding that supported this work.

## Authorship Contributions

*Participated in research design:* E.F. Douglass Jr., A.E. Ray

*Conducted experiments:* A.E. Ray, E.F. Douglass Jr.

*Performed data analysis:* A.E. Ray, E.F. Douglass Jr.

*Wrote or contributed to the writing of the manuscript:* A.E. Ray, E.F. Douglass Jr.

