## Supplementary material for "A Quantitative Approach for Assessing Multidrug Resistance in Cancer": Fig S1

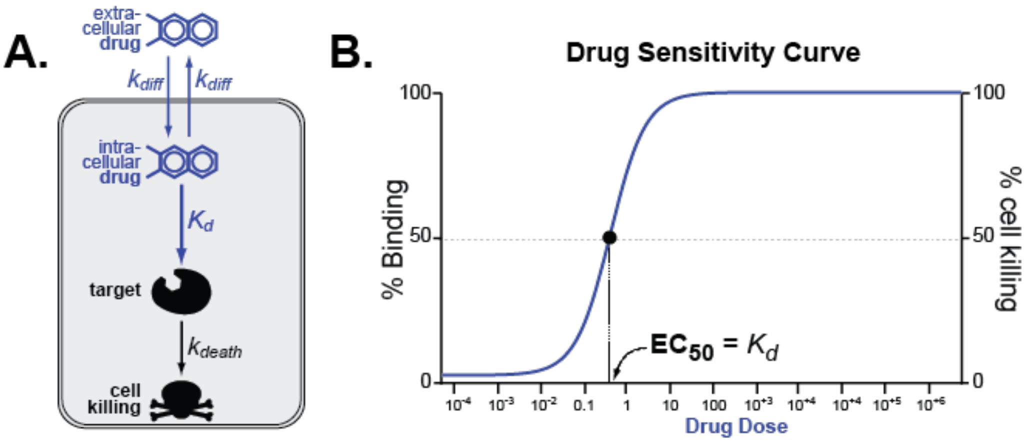


Fig S1. Mathematics of Drug Dose Response. The $EC_{50}$ can be approximated to the $K_d$ in normal tissues. (A) Drug diffuses into cells where the [drug]in equals the [drug]out. Drug binds to the intracellular target measured through the $K_d$. Bound drug causes cell death measured through the $k_{death}$. (B) The % binding to the intracellular target can be approximated to the % cell killing in the drug dose response curve.

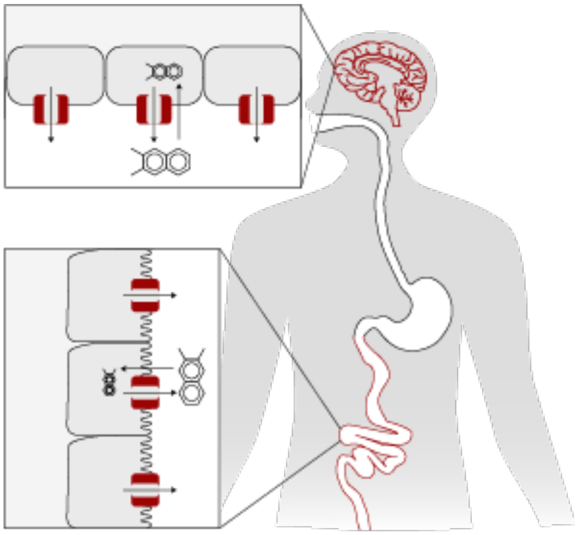

Fig S2. Pgp Normal Expression in Tissues. Pgp is typically expressed in tissues with pharmacokinetics and barrier functions such as the brain, intestines, liver and kidneys.

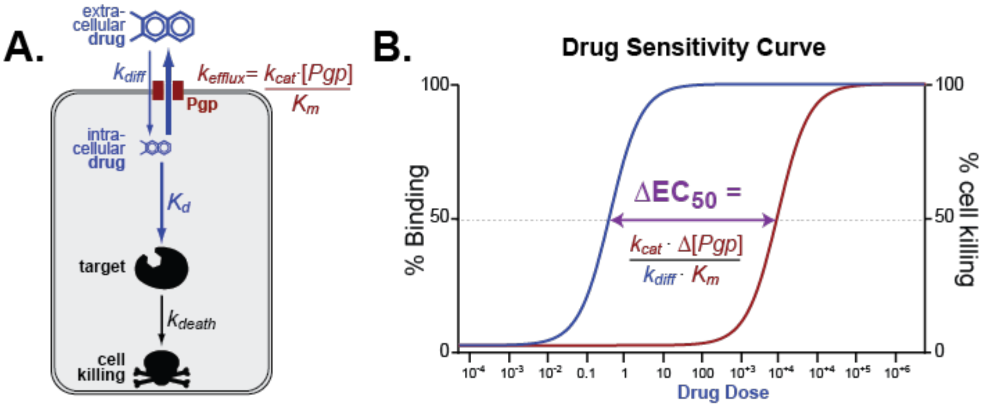


Fig S3. Drug Dose Response in MDR. Multidrug resistant cancer cells increase the $EC_{50}$. (A) Multidrug resistant cancer cells have increased Pgp expression. Pgp increases drug efflux, so [drug]out >> [drug]in measured through the $k_{efflux}$. Less drug is available to bind to the intracellular target and cause cancer cell death. (B) Pgp expression shifts the $EC_{50}$ right because a larger drug dose is required to achieve 50% binding and cell killing.

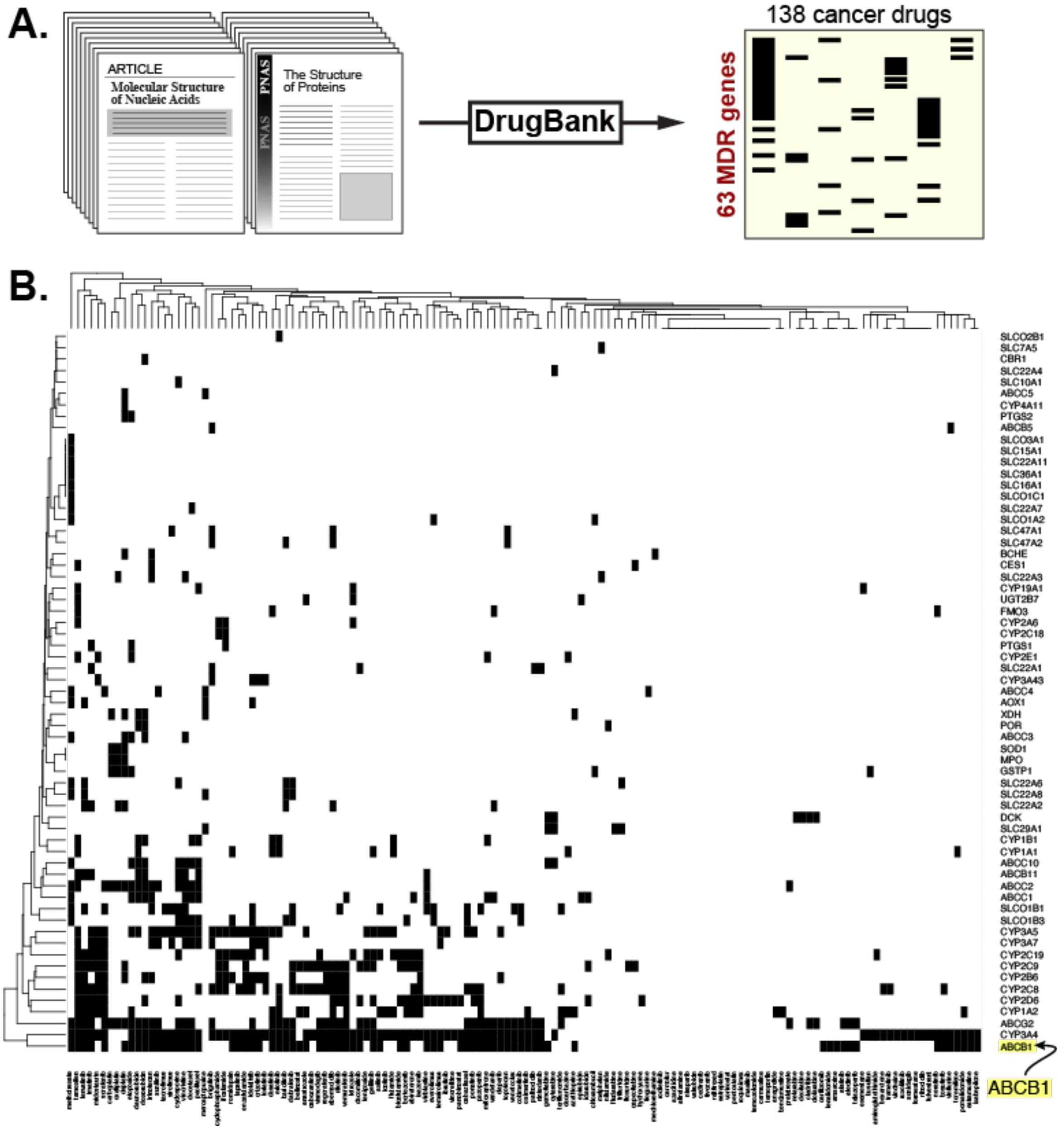


Fig S4. DrugBank MDR Genes and Drugs. DrugBank data on 63 MDR genes and 138 drugs. (A) DrugBank data is derived from the scientific literature. Sixty-three MDR genes representing ABC and SLC transporters and CYP450 metabolism were paired with 138 FDA approved oncology drugs. (B) Most current knowledge of MDR genes from the scientific literature is on ABCB1, the gene for Pgp.

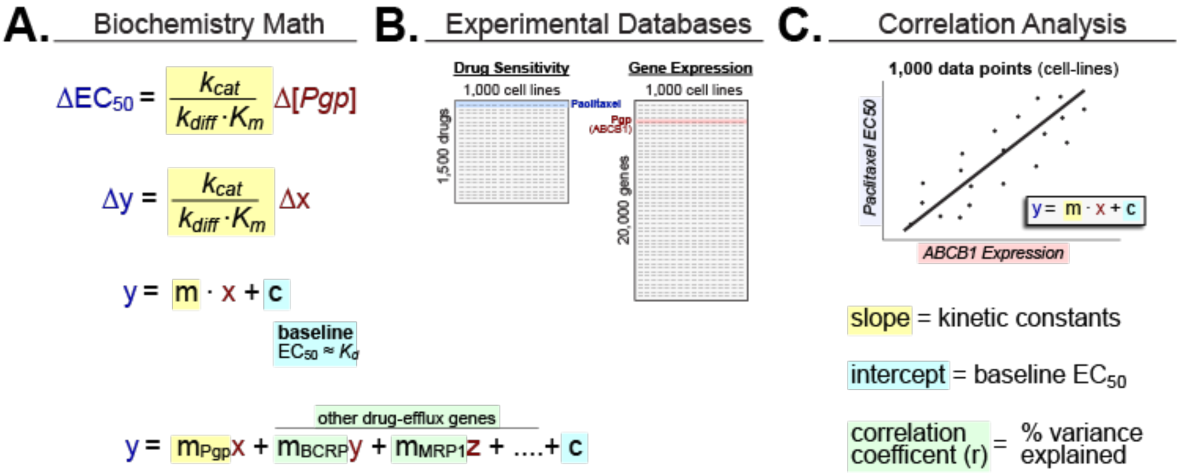


Figure S5. Mathematically Modeling MDR. The $EC_{50}$ shift, ABCB1 expression and enzyme kinetics can be represented through the linear equation, y = m * x. (A) Enzymes kinetics, $k_{cat}$ / ($k_{diff}$ * $K_m$), represent the slope (m) as it determines how y, the $EC_{50,}$ changes with x, ABCB1 expression. As a phenotype, MDR is represented by the independent contributions of several drug efflux genes. (B) CCLE has ABCB1 mRNA expression data across 1,000 cancer cell lines. PRISM has oncology drug $EC_{50}$ data across 479 of 1,000 cancer cell lines. (C) A correlation analysis compares CCLE ABCB1 mRNA expression with PRISM oncology drug $EC_{50}$ where single data points are individual cancer cell lines.

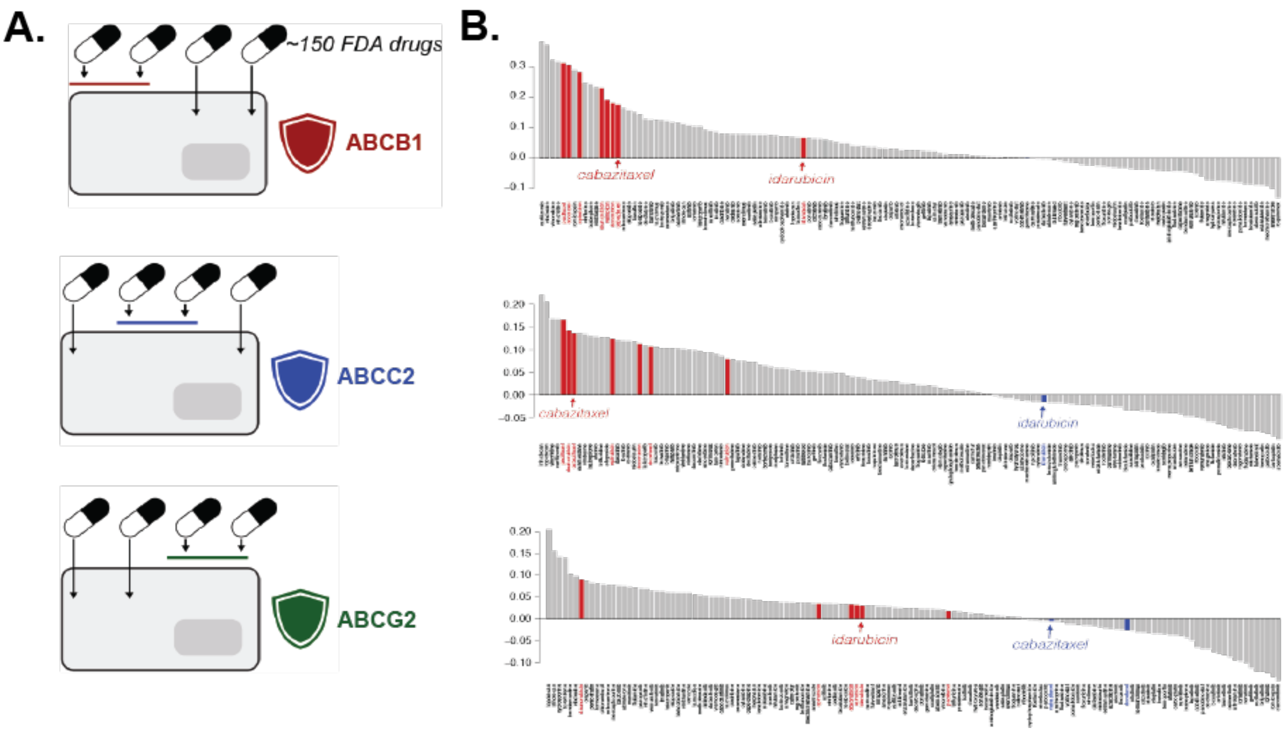


Fig S6. Quantitative Rankings of Drugs. Waterfall plots rank the FDA approved library of 138 oncology drugs by Pearson coefficient. (A) Different drugs are resisted by different armors such as transporters the cancer expresses to protect itself. (B) Higher to lower ranked drugs are more to less likely to be resisted by the cancer where Pearson coefficients indicate the strength of MDR substrate specificity.

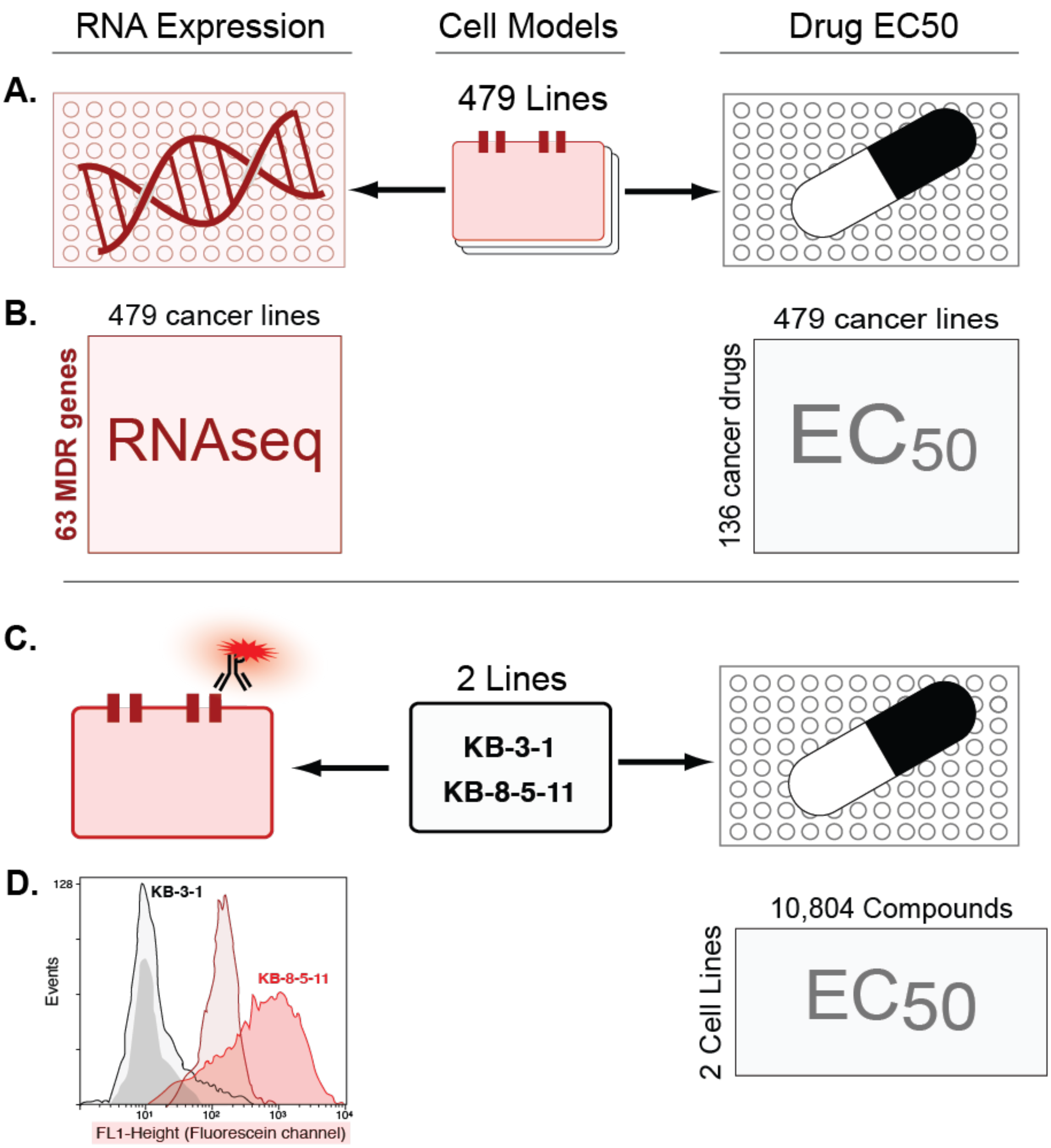


Fig S7. Approaches for Substrate Specificity Metrics. Quantitative metrics for Pgp substrate specificity are obtained through 2 approaches. (A) CCLE and PRISM databases characterize 479 cancer lines through mRNA expression and drug $EC_{50}$. (B) CCLE RNAseq analysis of 63 MDR genes and PRISM $EC_{50}$ analysis of 138 oncology drugs across 479 cancer lines. (C) KB-3-1 parent and KB-8-5-11 Pgp-high cancer lines characterized through protein expression and drug $EC_{50}$. (D) Flow cytometry analysis to quantify Pgp expression and drug screen of 10,804 compounds to assess Pgp substrate specificity.

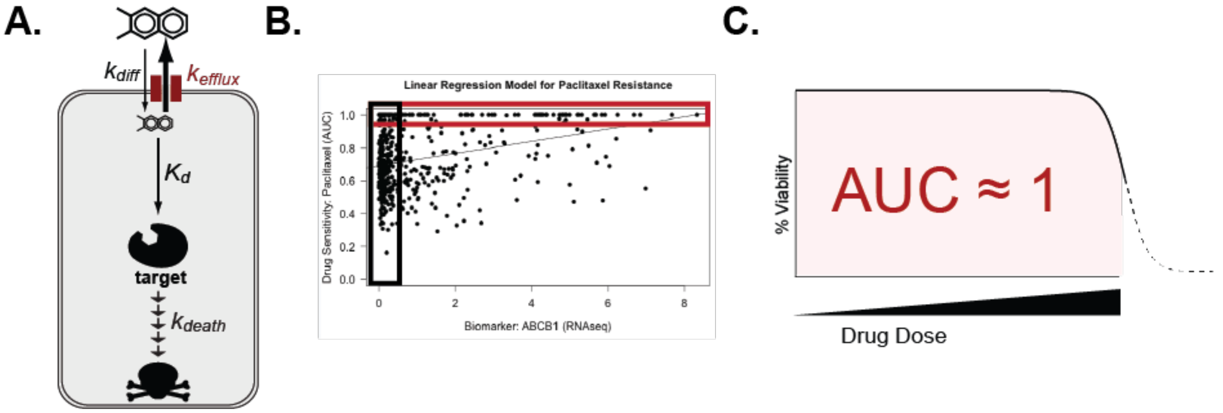


Fig S8. Limitations of AUC in Drug Screening. The AUC does not yield a measurable $EC_{50}$ without cytotoxicity. (A) For cytotoxicity to occur, drug must bind the intracellular target and cause cell death over time. (B) In our correlation analysis, there are vertical and horizontal clusters of cell lines (single data points). The vertical cluster includes drug sensitive cell lines which have low to no ABCB1 expression. The horizontal cluster includes drug resistant cell lines which do not exhibit cytotoxicity regardless of ABCB1 expression. (C) An AUC = 1 means that the drug does not cause cytotoxicity at the drug concentrations tested or the drug is not cytotoxic to the cell line regardless of concentration.

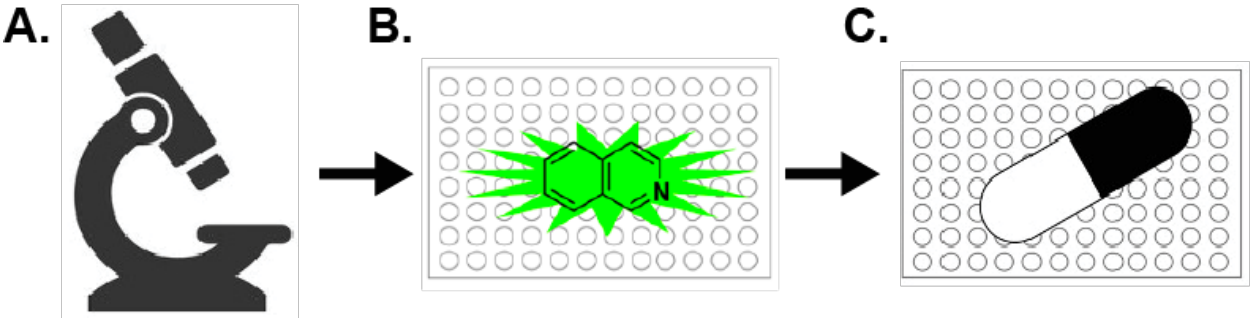


Fig S9. Experimental Pipeline for Assay Development. (A) Fluorescent microscopy to optimize cell culture and Calcein AM concentration. (B) Plate-based assay with Calcein AM to optimize competition with Pgp inhibitors. (C) Plate-based screen with Calcein AM to test 76 drugs from the FDA approved oncology drug library.
